# Spatial multi-omics reveal intratumoral humoral immunity niches associated with tertiary lymphoid structures in pancreatic cancer immunotherapy pathologic responders

**DOI:** 10.1101/2024.09.22.613714

**Authors:** Dimitrios N. Sidiropoulos, Sarah M. Shin, Meredith Wetzel, Alexander A. Girgis, Daniel Bergman, Ludmila Danilova, Susheel Perikala, Daniel H. Shu, Janelle M. Montagne, Atul Deshpande, James Leatherman, Lucie Dequiedt, Victoria Jacobs, Aleksandra Ogurtsova, Guanglan Mo, Xuan Yuan, Dmitrijs Lvovs, Genevieve Stein-O’Brien, Mark Yarchoan, Qingfeng Zhu, Elizabeth I. Harper, Ashani T. Weeraratna, Ashley L. Kiemen, Elizabeth M. Jaffee, Lei Zheng, Won Jin Ho, Robert A. Anders, Elana J. Fertig, Luciane T. Kagohara

## Abstract

Pancreatic adenocarcinoma (PDAC) is a rapidly progressing cancer that responds poorly to immunotherapies. Intratumoral tertiary lymphoid structures (TLS) have been associated with rare long-term PDAC survivors, but the role of TLS in PDAC and their spatial relationships within the context of the broader tumor microenvironment remain unknown. We generated a spatial multi-omics atlas encompassing 26 PDAC tumors from patients treated with combination immunotherapies. Using machine learning-enabled H&E image classification models and unsupervised gene expression matrix factorization methods for spatial transcriptomics, we characterized cellular states within TLS niches spanning across distinct morphologies and immunotherapies. Unsupervised learning generated a TLS-specific spatial gene expression signature that significantly associates with improved survival in PDAC patients. These analyses demonstrate TLS-associated intratumoral B cell maturation in pathological responders, confirmed with spatial proteomics and BCR profiling. Our study also identifies spatial features of pathologic immune responses, revealing TLS maturation colocalizing with IgG/IgA distribution and extracellular matrix remodeling.

**GRAPHICAL ABSTRACT:** 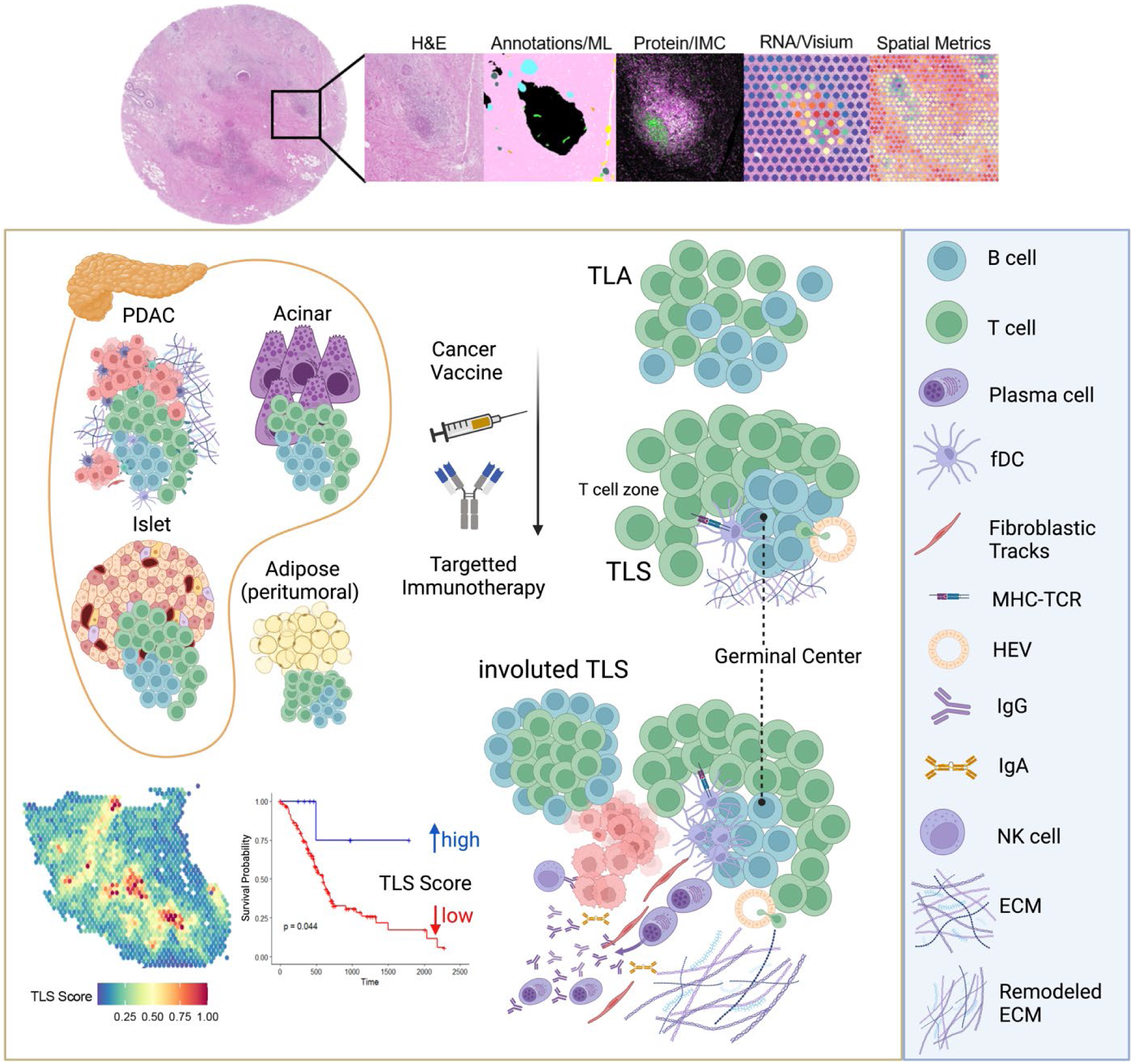

**HIGHLIGHTS:**

- Integrated multi-modal spatial profiling of human PDAC tumors from neoadjuvant immunotherapy clinical trials reveal diverse spatial niches enriched in TLS.
- TLS maturity is influenced by tumor location and the cellular neighborhoods in which TLS immune cells are recruited.
- Unsupervised machine learning of genome-wide signatures on spatial transcriptomics data characterizes the TLS-enriched TME and associates TLS transcriptomes with survival outcomes in PDAC.
- Interactions of spatially variable gene expression patterns showed TLS maturation is coupled with immunoglobulin distribution and ECM remodeling in pathologic responders.
- Intratumoral plasma cell and immunoglobin gene expression spatial dynamics demonstrate trafficking of TLS-driven humoral immunity in the PDAC TME.

**Significance:** We report a spatial multi-omics atlas of PDAC tumors from a series of immunotherapy neoadjuvant clinical trials. Intratumorally, pathologic responders exhibit mature TLS that propagate plasma cells into malignant niches. Our findings offer insights on the role of TLS-associated humoral immunity and stromal remodeling during immunotherapy treatment.

## INTRODUCTION

Cancer immunotherapies provide significant clinical benefit for patients with some cancers, generally those that present with baseline immunogenicity (Chan et al., 2019; Yarchoan et al., 2017). Non-immunogenic solid tumors, however, such as pancreatic ductal adenocarcinoma (PDAC), are generally characterized by a tumor microenvironment (TME) that is resistant to immunotherapies including immune checkpoint blockade (Timmer et al., 2021). This is mostly attributed to a low tumor mutational burden, low immunogenicity, a highly desmoplastic stroma, and immunosuppressive cellular populations within the TME (Maleki Vareki, 2018). Because of its poor response to chemotherapy, radiotherapy, and immunotherapy, it is projected that PDAC will become the second leading cause of cancer-related deaths in the US by 2030 (Siegel et al., 2019).

In immunotherapy-responsive cancers that are typically immune infiltrated such as melanoma, tertiary lymphoid structures (TLS) have been described as tissue-embedded organized immune cellular neighborhoods (Cabrita et al., 2020). Regardless of treatment, the presence of TLS within the TME is associated with increased patient survival (Sautès-Fridman et al., 2019). Studies of TLS have shown that TLS development undergoes different stages of maturation that culminates in the formation of germinal centers and dissemination B cells, including plasma cells (Teillaud et al., 2024). In aPD1/aPDL1 immune checkpoint blockade therapy trials, increased density of plasma cells are associated with more favorable outcomes in several cancers, including non-small cell lung cancer (Patil et al., 2022), and soft-tissue sarcomas with TLS (Italiano et al., 2022).

In PDAC, rare, activated TLS presence in untreated patients is correlated with improved survival rates, with studies indicating a significant survival benefit in patients with mature TLS (Castino et al., 2016; Gunderson et al., 2021). These findings support that inducing TLS and applying TLS assessment in clinical practice could enhance and guide therapeutic strategies (Petroni et al., 2024). We previously reported the induction of intratumoral TLS following administration of a neoadjuvant GM-CSF-secreting allogeneic vaccine (GVAX) to PDAC patients in combination with low dose cyclophosphamide (NCT00727441) (Lutz et al., 2014). However, no clinical benefit was observed, likely owing to immune tolerance and immunosuppressive mechanisms in the PDAC TME. It was hypothesized that combination immunotherapy strategies may overcome these limitations and further augment positive pathological responses to induce significant survival benefit. In subsequent clinical trials in PDAC combining GVAX with immune checkpoint blockade and 41BB agonism in the neoadjuvant setting presented with promising clinical and pathologic responses (NCT02451982) (Heumann et al., 2023; Muth et al., 2021). We hypothesized that these combination strategies may have also affected TLS maturation in a synergistic manner enabling cellular crosstalk intratumorally. Specifically, we expected that favorable effects of this crosstalk enhance recruitment and activation of immune cells, such as CD8^+^ T cells and B cells, facilitated by chemokines and cytokines within the TME. However, the relative differences in maturation and function between TLS in different tumor regions may be attributed to several factors, including higher density of immune cells, antigenic targets, and stromal elements supporting prolonged and effective immune responses. Understanding these mechanisms should also lead to improved therapeutic approaches that enhance TLS maturation and prolong patient survival.

Spatial molecular characterization of TLS in the PDAC TME during immunotherapy treatment can elucidate the impact of the broader TME on their function. In this study, we generated integrated cellular and molecular maps of the TLS-enriched human PDAC TME from biospecimens taken from our unique patient cohorts treated in our neoadjuvant immunotherapy clinical trials. Comparing distinct therapeutic combinations tested in our novel platform neo-adjuvant clinical trial with patient outcomes provides the potential to distinguish the cellular and molecular landscapes in the TME that impact TLS maturation. Specifically, we performed spatial transcriptomics and spatial proteomics analysis of TLS-enriched PDAC TME as well as tumor-adjacent lymph nodes from patient tumors following treatment with GVAX and the antagonist anti-PD-1 antibody nivolumab (aPD1) alone or in combination with the agonist anti-41BB antibody urelumab (a41BB). Applying machine learning methods to these data generated gene expression signatures of TLS maturation. These signatures distinguished patients with longer survival in TCGA and further enabled us to uncover activated B cell transcriptional programs emerging from mature TLS niches colocalizing with stromal remodeling. Overall, our work sheds light on the TME of immunotherapy pathologic responders and contributes spatially defined transcriptional markers of TLS in PDACs.

## RESULTS

### Spatial distributions of tertiary lymphoid structures highlight distinct intratumoral niches in PDAC treated with neoadjuvant immunotherapies

Previous studies of human PDAC tumors have shown that scoring patient tumors by TLS density and location showed that rare long-term survivors (<2%) present high intratumoral density of TLS, whereas TLS in typical short-term survivors’ tumors are limited and peritumoral (Hiraoka et al., 2015). TLS location is a critical feature in deploying anti-tumor immunity, yet an in-depth characterization of intratumoral TLS spatial niches in PDAC tumors has been lacking. To better understand the spatial organization and functional implications of lymphoid aggregates within the PDAC TME, we aimed to conduct an integrated histological, as well as proteomic and transcriptomic spatial analyses of PDAC patient tumor samples surgically resected after neoadjuvant immunotherapy combinations clinical trials (Fig. 1A). For this purpose, we collected frozen and formalin-fixed, and paraffin embedded (FFPE) samples from 26 PDAC patients treated with a combination of GVAX, anti-PD1 (nivolumab), 41BB agonist (urelumab) and cyclophosphamide (Cy). Previous pathological studies of the biospecimens from these trials demonstrated that a subset of tumors exhibited pathologic responses and intratumoral lymphoid aggregate induction (Heumann et al., 2023; Lutz et al., 2014). In this study, we performed spatial multi-omics profiling on a cohort of samples selected for TLS enrichment, including both pathological responders and non-responders across treatment conditions as well as matched lymph nodes to enable comprehensive profiling of germinal center maturation in TLS and lymph nodes treated with immunotherapy (Fig. 1B).

**Figure 1.**
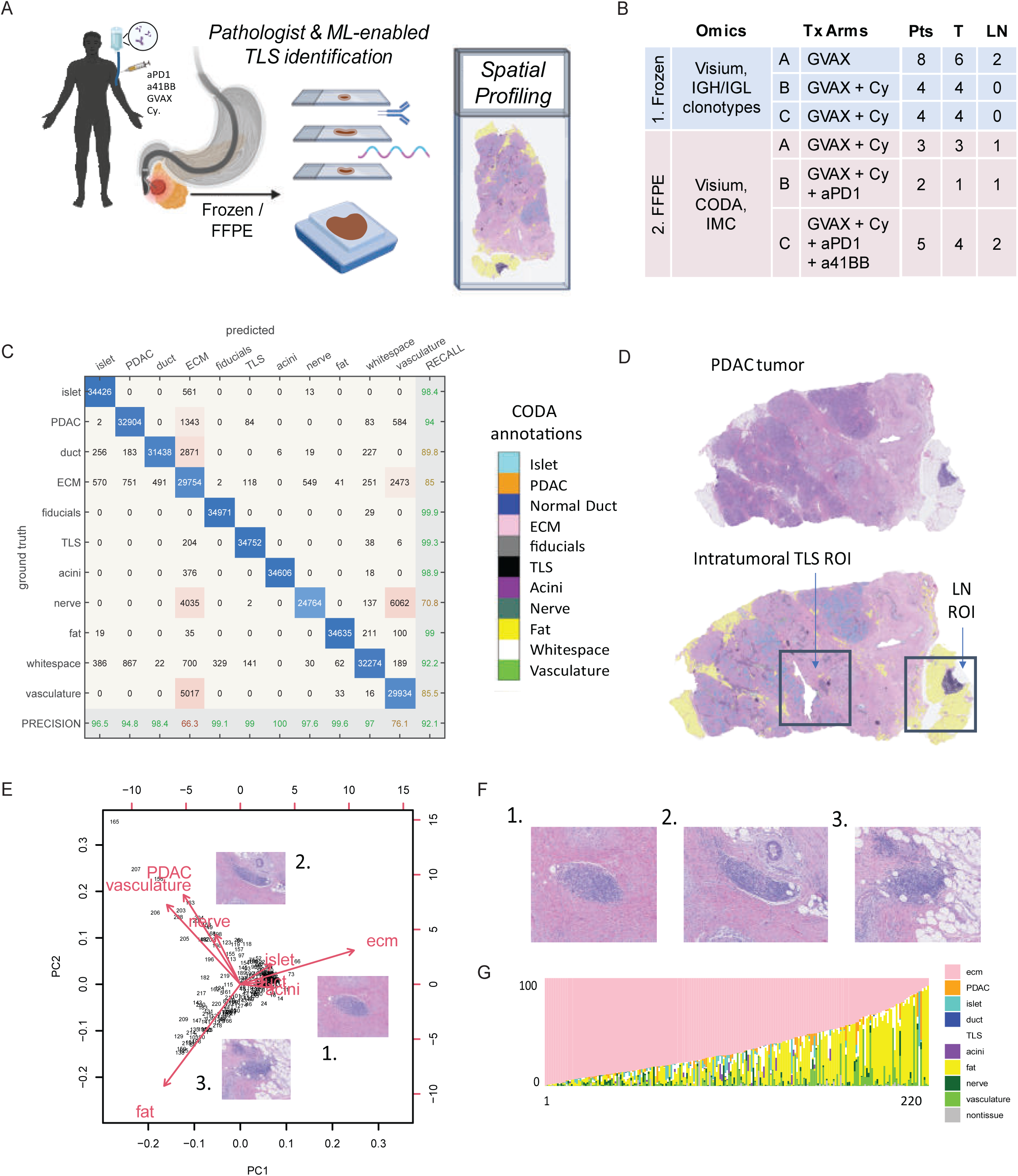
Leveraging a neoadjuvant immunotherapy clinical trial platform and spatial omics of resected pancreatic tumors to elucidate TLS in PDAC. **A**. Study design of lymphoid organ spatial omics profiling in neoadjuvant immunotherapy treated PDAC. **B.** Metadata for samples used in the generation of the spatial omics in this study. **C**. Plot showing CODA model performance. Shown diagonally are the total number of pixels correctly predicted by CODA in the testing set when compared to pathology annotations. Precision per class as a percentage of correctly annotated pixels is shown in gray column. **D**. Representative image of a PDAC tumor H&E from FFPE cohort (patient pub ID P_45) before and after pixel classification with CODA. **E**. PCA plot showing PC1 and PC2 of TLS niche compositions by CODA class, highlighting three putative TLS niche subclasses (**F**). **G**. Stacked barplot of 220 predicted TLS showing tissue composition of each of their niches ordered by decreasing ECM/stroma composition predicted by CODA.

For each sample, histological examination was performed by an expert pathologist to establish tissue identity and pathology. In FFPE samples, pathologist annotations were also used to train a machine learning-enabled H&E segmentation model using CODA (Kiemen et al., 2022) to label at the pixel level histological classes of interest, including malignant ducts, normal ducts, extracellular matrix components, immune cells, acinar cells and adipose tissue (Fig. 1C). Each of these classes extrapolated pathology annotations and segmented pixels with an overall model accuracy of 92.1% on an independent testing dataset. The trained model allowed us to identify lymphoid aggregates, putative TLS, and other tissue structures from whole slide H&Es (Fig. 1D).

The annotation of cellular features from H&E further enabled us to extract their neighborhood spatial compositions. For this analysis, tiles measuring 100x100nm^2^ were centered around each putative TLS (n=220). Pixel-level classifications were extracted from each tile to compute the percentage of pixels belonging to each of the learned classes to estimate spatial niche compositions per tile. We sought to classify the variation in the cellular neighborhoods in which TLS occur with a principal component analysis on the class composition matrix (Fig. 1E). A class of putative TLS are embedded in adipose tissue, consistent with peritumoral lymphoid structures. Characterization of intratumoral TLS niches demonstrated that intratumoral TLS are observed in a range of ductal, endocrine, exocrine, or stromal components (Figs. 1F, 1G). Although the distinction between peritumoral and intratumoral TLS have been previously described and studied in terms of their divergent impact on the PDAC TME, a thorough characterization of distinct intratumoral niches and the cellular crosstalk between immune cells within TLS and surrounding cells remains to be elucidated.

### Imaging mass cytometry reconstructs TLS lifecycle stages at single cell resolution revealing improved TLS maturation and function in pathologic responders

TLS development and maturation have been associated with the impact of the immune response during immunotherapy treatment (Teillaud et al., 2024). The heterogeneity of putative TLS tissue neighborhoods in our cohort prompted us to explore whether lymphoid aggregate-mediated structure formation and immunity specifically target and remodel local tissue contexture. Although our machine learning model trained on H&E can automatically identify immune dense regions suggestive of TLS, more refined immune cell characterization is required for definitive classification of TLS. Since H&E-stained tumor sections alone do not lend information about the cellular components of TLS, we applied Imaging Mass Cytometry (IMC) using a custom TLS panel of 35 markers (Supplemental Table S1) to confirm cell composition and assess structure and functional markers at single cell resolution to adjacent sections of those analyzed with the CODA annotation of H&E imaging. Following staining, representative 1mm^2^ tiles of from each tumor specimen were sampled by pathologist and ablated. The acquired images of protein targets were captured and used to resolve predicted TLS niches (n=17 tiles) reported by CODA as well as tumor adjacent lymph nodes (n=6 tiles) (Supplemental Data S1).

Pearson’s correlation of IMC protein markers in segmented single cells in tiles containing intratumoral TLS of responders versus nonresponders suggested diverging spatial distributions of immune cell types, including B cell subtype markers (CD20, CD138, CD23, CXCR3, AID) (Supplemental Figs. S1A, S1B). IMC tiles were examined for the presence or lack of structure in intratumoral TLS of both pathologic responders and nonresponders. The spatial distribution of immune cells within a TLS is commonly called zoning, and is understood as an important indicator of mature and functional TLS, along with the presence of a germinal center and high endothelial venules (HEV) (Fridman et al., 2023). Thus, T cell (CD3^+^) and B cell (CD20^+^) compartmentalization, presence of follicular dendritic cells (fDC, CD21^+^ cells) and PNAD^+^ endothelial cells were used as indicators of maturity in IMC tiles with putative TLS. TLS lacking spatially separated T/B cell compartments containing diffuse HEV and follicular dendritic cells (fDC) clusters were identified in acinar cell and islet-rich niches of pathological nonresponders (Figs. 2A, 2B). Notably, in a pathological responder to GVAX and nivolumab, we found two distinct TLS morphologies in a region of a resolving necrotic duct infiltrated with lymphoctes (Figs. 2C, 2D). Mature TLS presented with a densely clustered singular germinal center surrounded by T cells (Fig. 2E), whereas a secondary TLS with an involuted morphology was characterized by B cell compartments surrounding a T cell zone and disorganized fDC (Fig. 2F, 2G). Both morphologies were enriched in cells expressing cytotoxic marker GZMB, memory marker CD45RO and late-differentiated effector marker CD57. TLS were identified proximal to a resolving necrotic duct infiltrated with lymphocytes.

**Figure 2.**
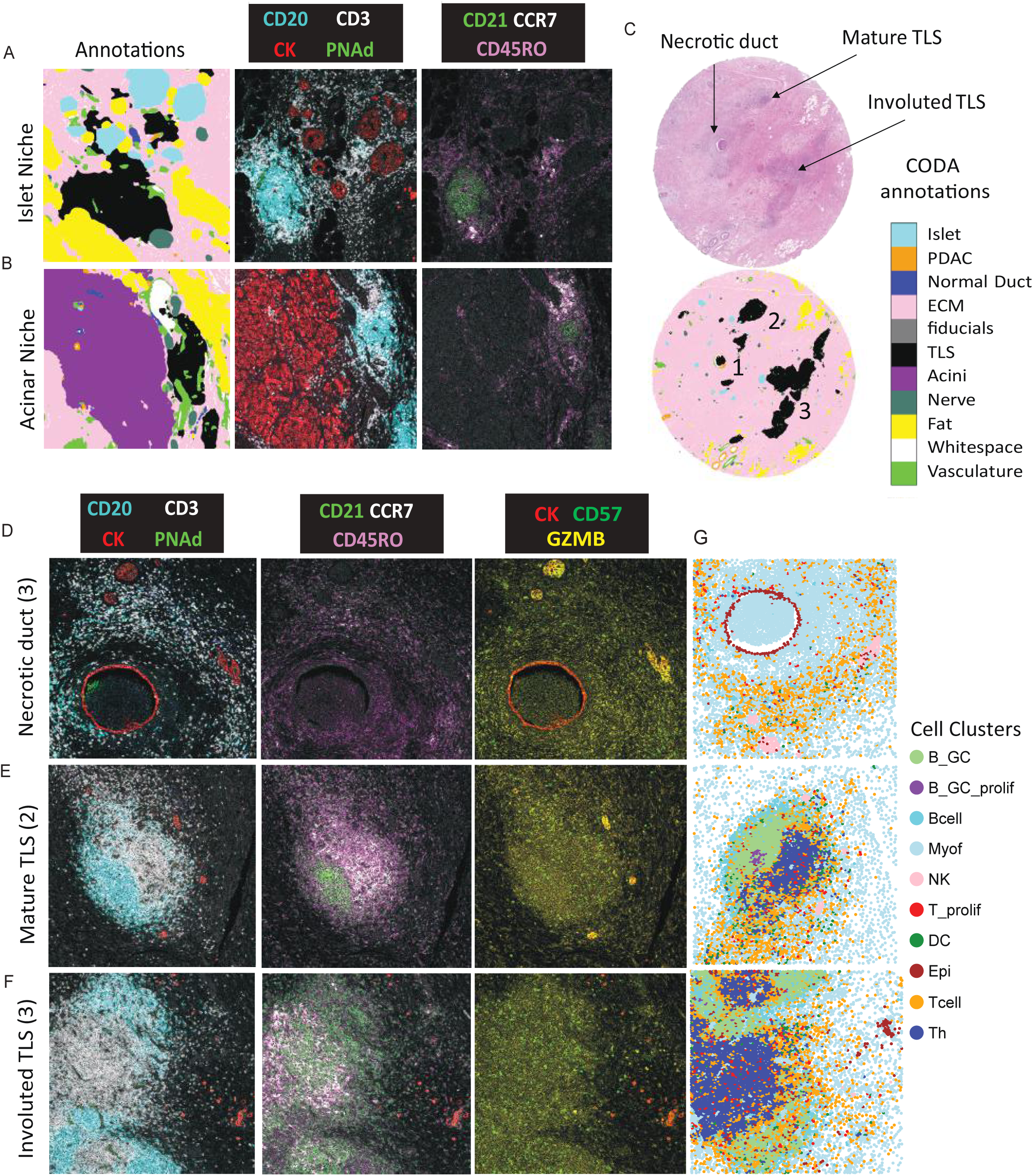
Resolving putative TLS and their cellular niches at single cell resolution using imaging mass cytometry (IMC). Pathology annotations and CD20 (blue), CK (red), CD3 (white), PNAd (green) as well as CD21 (green), CD45RO (pink), CCR7 (white) expression in an islet (**A**) and acinar (**B**) niche. **C**. Pathology annotations in pathologic responder, highlighting three regions: necrotic duct (1), mature TLS (2), involuted TLS (3). Three columns of panels showing expression of CD20 (blue), CK (red), CD3 (white), and PNAd (green); CD21 (green), CD45RO (pink), CCR7 (white); GZMB (yellow), CK (red) and CD57 (darkgreen) in each region respectively (**D-F**). Morphologies of necrotic duct (**G**), mature TLS (**H**) and involuted TLS (**I**) are illustrated using single cell clusters after image segmentation and clustering.

To further examine the cell types associated with distinct TLS niches and pathological response, we utilized the IMC data to compare the immune composition analysis between sample types. IMC imaging data were segmented into single cells. Unsupervised clustering was applied on protein expression profiles of segmented cells to annotate cell types. A total of 10 distinct cell clusters were annotated based on canonical cell type marker expression and location (Supplemental Figs. S1C, S2A). These include germinal center (GC) associated B and T cell subtypes, as well as dendritic cells (DC, DCSIGN+), NK cells (CD3^-^, CD57^+^, GZMB^+^), and pan-epithelial cells (CK^+^) and pan-fibroblasts (SMA^+^, PDPN^+^).

To evaluate immune cell function in the TLS-driven pathological response, differential cell type abundance, nearest neighbor and protein marker expression analyses were performed across all clustered IMC cell types comparing tiles from pathological responders (n=17) to nonresponders (n=6) (Supplemental Data S1). Cell type abundance and nearest cell neighbor composition profiles demonstrate higher density of germinal center B cells (B_GC) and T helper (Th) cells forming in intratumoral TLS and lymph nodes of responders (Supplemental Figs. S2B, S2C, S3A, S3B). In GC B cells, antigen presentation gene HLADR (FDR adj. p<0.001, Wilcoxon), proliferation marker KI67 (FDR adj. p<0.05, Wilcoxon) and trafficking markers CXCR3 and CD138 (FDR adj. p<0.001, Wilcoxon) were significantly increased in responders. However, detectable expression of the plasma cell marker CD138 was only observed in responder intratumoral TLS samples and not in nonresponders (Supplemental Figs 1A,1B; Supplemental Data S2). In Th cells, naive marker CD45RA was significantly higher (FDR adj. p=0.0004, Wilcoxon) in responders whereas memory marker CD45RO was not significantly different. In the pan-fibroblast cluster, higher expression of HLADR, PDPN, CXCR3 and CXCR5 (FDR adj. p<0.001, Wilcoxon) were also observed in nonresponders, suggesting antigen presenting CAF subtypes and CAF-mediated immune recruitment or retention in TLS niches. Neighborhood composition analysis showed T cell clusters as the top non-self-neighbors of fibroblasts, followed by Epithelial, B and NK cells (Supplemental Figs. S3A, S3B). These T cells were comprised of a mixture of Th (CD4) and Tc (CD8) subpopulations but lacked expression of FOXP3 or other immunosuppressive markers (Supplemental Figs. S1C).

### Mapping TLS transcriptomes in the PDAC tumor microenvironment

Image IMC resolved the immune compartments of TLS in targeted regions at single cell resolution. Given the known complexity of the PDAC TME, our next aim was to associate specific histological structures with gene expression patterns to infer molecular mechanisms in TLS from different cellular neighborhoods using spatial transcriptomics. TLS resemble lymph nodes (LN) in their organization and function, comprising similar cellular components like B cells, T cells, and follicular dendritic cells that orchestrate immune responses (Sautès-Fridman et al., 2019). Thus, we included both TLS and LN samples in spatial transcriptomics experiments to understand how different TLS niches mirror or diverge from LNs in their role in orchestrating tumor immunity during neoadjuvant immunotherapy in PDAC patients. Specifically, we profiled tumors and associated tumor-adjacent lymph nodes from 26 patients with PDAC treated with combination neoadjuvant immunotherapies (GVAX n=19, GVAX+aPD1 n=2, GVAX+aPD1+a41BB n=5). In total, we generated spatial transcriptomics (ST) data from 28 Visium individual capture areas, including 22 tumors and 6 tumor associated lymph nodes from frozen (n=16, Fig. 3A) and FFPE (n=12, Fig. 3B) samples. To map individual histological features to gene expression in the ST data, we labeled individual Visium spots by pathology annotations based on the matched H&E imaging data. On frozen samples, we manually transferred expert PDAC pathologist tissue annotations onto the ST data. In FFPE samples, since tissue morphology is fixed and retained, CODA was used to assist in extrapolating pathologist annotations across all pixels of the ST images (Fig. 3B, Supplemental Fig. S4) (Kiemen et al., 2022; Bell et al., 2024). Visium spots are 55um wide, thus capturing more than one cell resulting in mixed histological and transcriptional signatures. Hence, we trained a nuclei detection model and annotated ST spots after accounting for the CODA H&E annotation class with the highest proportion of annotated H&E image pixels and annotated nuclei per spot (Supplemental Fig. S4A-C). Unsurprisingly, the tissue annotation classes with the highest number of nuclei were TLS (Supplemental Fig. S4D).

**Figure 3.**
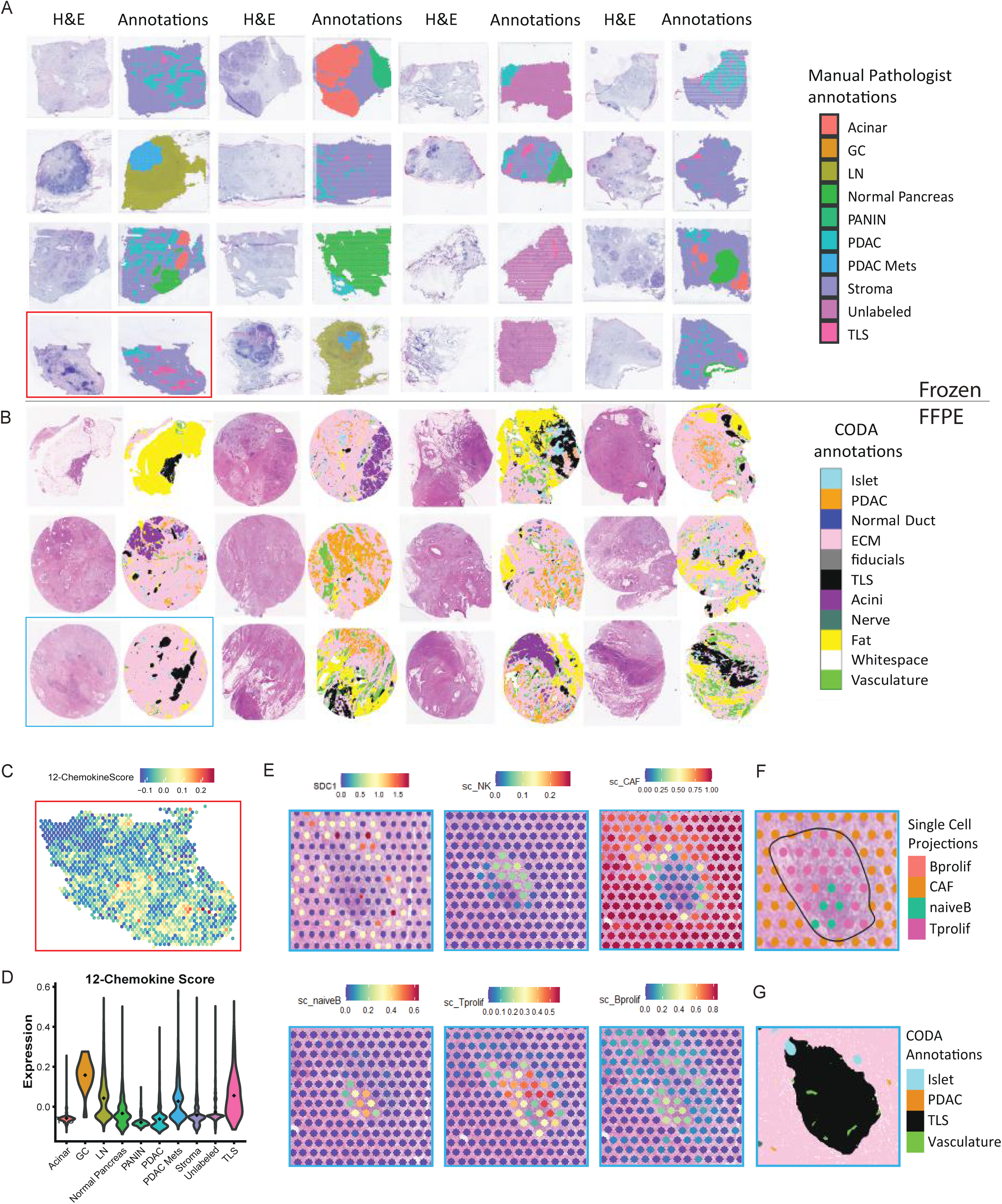
Spatial Transcriptomics atlas of the PDAC TME treated with neoadjuvant immunotherapies incorporating two clinical cohorts and archival tissue methodologies. **A**. Spatial transcriptomics of 14 frozen PDAC tumors and 2 adjacent lymph nodes. **B**. Spatial transcriptomics of 12 FFPE samples from 8 PDAC tumors and 4 adjacent lymph nodes. For each sample, H&E and pathology annotations are shown. For frozen samples, manual pathology annotations are illustrated (A), whereas in FFPE samples, pathologist trained CODA H&E pixel classifier annotations are shown (B). **C**. 12-chemokine score expressed on a representative sample with high TLS density, boxed in red. **D**. Expression of 12-chemokine score across pathology annotations in the frozen cohort. **E.** Expression of CXCL13 highlighting mature TLS in sample boxed in blue, as well as single cell deconvolution results illustrating density of NK cells, CAFs, naïve B cells, proliferating T cells and B cells. **F.** Top cell type from deconvolution in representative mature TLS and corresponding pathology annotations from CODA (**G**).

To further confirm the TLS spots annotated from the H&E at the gene expression level, we aggregated the expression of a 12-chemokine signature and compared the resulting scores between spots with different cellular annotations (Coppola et al., 2011). This score along with other singular T cell and B cell markers (*CDE, CD8A, CD4, CD79A*) were highest in TLS and GC of LN compared to remaining histology classes (Fig. 3 C-D). While the H&E imaging gives broad annotation of tissue components, to infer the cellular and molecular changes in TLS we sought to further analyze the expression profiles for individual cell types and states within the same spot of interest using spot deconvolution. Integration anchors were computed in Seurat between the ST data and a curated PDAC single cell RNAseq reference dataset (Guinn al., 2024), which included annotations of cancer, fibroblast, lymphoid and myeloid cell types and subtypes collected from aggregated single cell RNA seq data of previously published PDAC studies. Spot deconvolution using Seurat projections of single cell clusters reported by Guinn et al. highlighted the spatial distribution of cell types found in and around TLS in the PDAC TME. Distinct T and B cell compartments intersecting with cancer associated fibroblasts (CAF) at the TLS border were observed in spots overlapping with mature TLS, suggesting crosstalk between the TLS and ECM in the stromal compartment (Fig. 3 E-F).

### Unsupervised machine learning identifies genome-wide spatial transcriptional signatures correlating with pathology annotations and disease outcomes in PDAC

Unlike single cell suspension omics, spatial omics aim to extract molecular information without affecting structure, potentially delivering novel expression profiles. Due to its unique ability to learn cellular states and further extensions to determine molecular changes from cell-cell interactions across niches, we applied our Bayesian non-negative matrix factorization (NMF) approach CoGAPS (Fertig et al., 2010) to on the ST data learn distinct coregulated gene expression patterns that specifically identify cellular communities in PDAC tissue including TLS (Deshpande et al., 2023). This implementation resulted in distinct patterns, each associated with a unique cluster of genes and tissue annotation, including validated TLS (Fig. 4A). In the frozen cohort where total mRNA transcriptome profiling is possible, we learned 10 patterns, of which the TLS pattern (Pattern 9) was most highly expressed in lymph node germinal centers (Figs. 4 B-D). This supports that TLS regions with higher values of this pattern were more GC-like, and potentially more mature which would indicate more organized, diverse, and active immune substructures. We sought to further annotate the genes associated with each pattern to determine their function by applying PatternMarkers to rank the set of genes that are uniquely associated with each pattern (Stein-O’Brien et al., 2017). As evidence of the relevance of the molecular weights of cells in TLS patterns, their set of pattern marker genes included a collection of 304 genes (Fig. 4A), many of which are immune related genes, including FOXP3, CCR7 and LTA (Supplemental Data S3).

**Figure 4.**
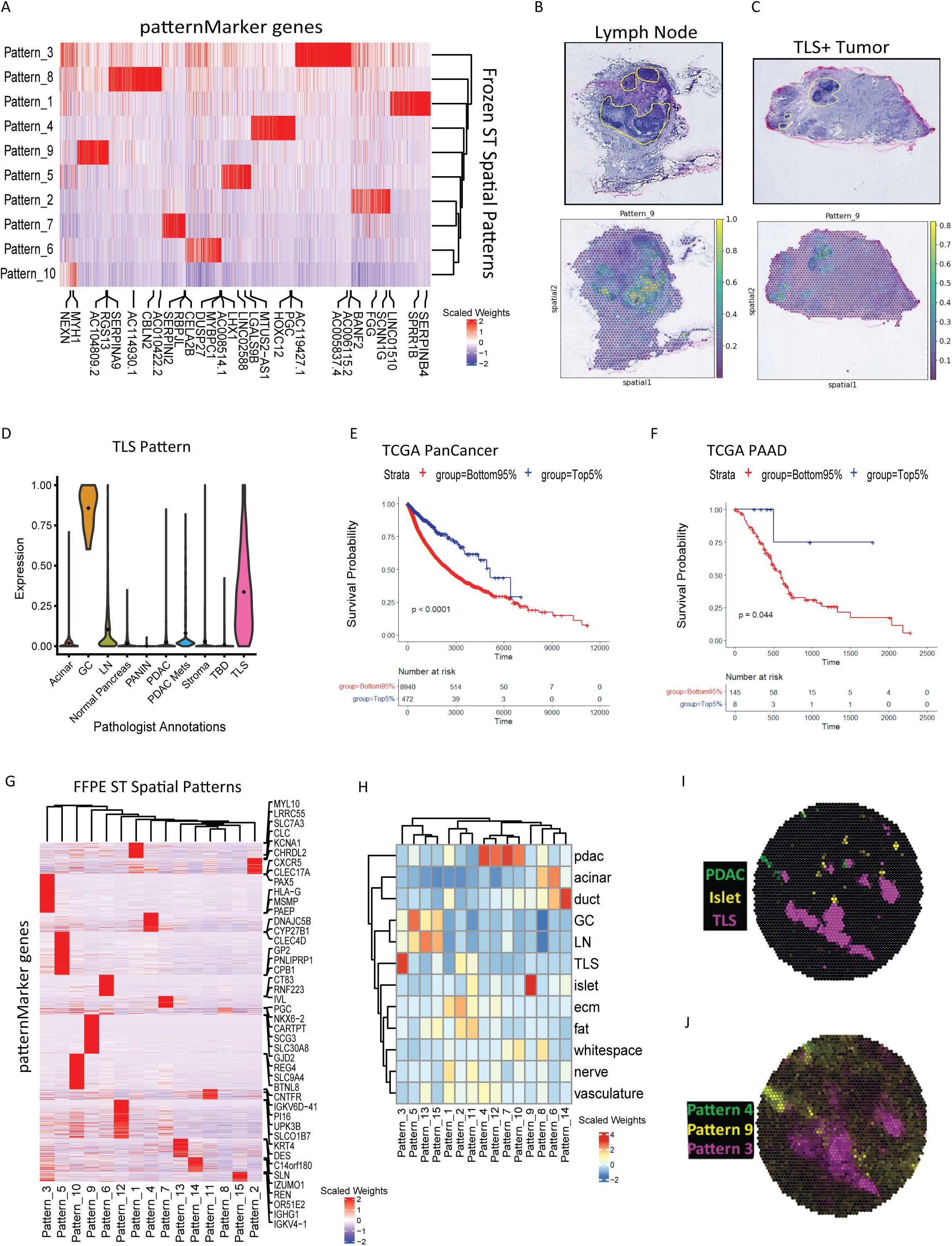
Matrix factorization resolves the spatial gene expression landscape of TLS enriched TME and tumor draining lymph nodes in PDAC and identifies signatures positively correlating with pathology annotation and disease outcomes. **A**. Heatmap showing top patternMarker genes and 10 spatial patterns in frozen cohort ST data. **B**. TLS pattern score (Pattern 9) expressed on a lymph node and a tumor sample enriched with TLS (**C**). **D**. TLS pattern score expressed across pathology annotations. Kaplan Meier curves of top 5^th^ percentile vs. bottom 95^th^ percentile of patients enriched with Pattern 9 projection scores in all cancer subtypes combined (**E,** p<0.001) and in PAAD alone (**F**, p =0.044). **G**. Heatmap showing top patternMarker genes and 15 spatial patterns in FFPE cohort ST data. **H**. Heatmap showing weights from each pattern aggregated by CODA annotations. **I**. Spatial ST plots showing CODA annotations of PDAC (green), Islets (yellow) and TLS (purple) as well as corresponding spatial patterns 4, 9 and 3 (**J**) in a representative pathologic responder sample.

Although TLS-related transcriptional signatures have been associated with more positive outcomes in a variety of cancers (Fridman et al., 2023), association of transcriptomic signatures mined directly from spatial transcriptomic data to outcomes has not been demonstrated in PDAC. To evaluate the clinical relevance of these patterns, we used our projectR transfer learning method to project each of them across bulk RNAseq data from The Cancer Genome Atlas (TCGA) (Sharma et al., 2020; Weinstein et al., 2013). We first projected each pattern to bulk RNAseq data pan-cancer and then computed a univariate Cox proportional hazards regression analysis to examine the impact of each pattern on overall survival. Pan-cancer, the TLS pattern had the most significant negative regression coefficient (p=1.4e-29) (Supplemental Fig. S5). In a Kaplan-Meier analysis, the top 5^th^ percentile of patients expressing TLS-pattern patternMarker genes showed significantly longer overall survival both pancancer (p<0.0001, Fig. 4E), and in PDAC-only subjects (p=0.044) (Fig. 4F). The transfer learning results further demonstrate the clinical relevance of the TLS spatial transcriptomic patterns and suggest that TLS patternMarker genes could potentially be used as surrogate markers for TLS in bulk RNAseq data.

### Intratumoral TLS niches coordinate local antigen sampling, immunoglobulin production, complement activation, and extracellular matrix remodeling

Given the association of high TLS signature enrichment with longer survival, we hypothesized that intratumoral TLS are propagating anti-tumor humoral immunity initiating a cascade of antibody opsonization and tumor cell clearance. To understand the roles these TLS have in the TME and the mechanisms with which they might promote anti-tumor immunity, we modeled the spatial dynamics of the gene expression patterns we discovered. We leveraged the CODA-annotated FFPE spatial transcriptomics dataset of the TLS to further determine the transcriptional changes associated with immune cell function in the stroma residing TLS relative to intratumoral TLS. Recently, we developed a computational approach called SpaceMarkers that characterizes the molecular changes from cell-cell interactions based on distinct NMF spatial patterns of interest (Deshpande et al., 2023). To map the molecular changes in immune cell hotspots that arise from TLS in distinct spatial niches in the pancreas, we applied CoGAPS in the FFPE cohort and learned 15 patterns correlated with CODA annotations (Figs. 4 G-J). Then using SpaceMarkers, we computed hotspots that define regions in which pattern pairs of interest are co-localized (Fig. 5A). The differentially expressed genes identified in overlapping and non-overlapping hotspots are reflective of the molecular changes associated with their interactions. Specifically, we looked at the genes differentially expressed at the intersection between TLS, stroma, pancreatic islet, and PDAC to identify associated gene expression patterns (Figs. 5A). Gene set enrichment of the differentially expressed genes at the intersection of TLS and stromal spatial patterns showed enrichment of humoral immunity pathways, complement activation, and cilium dependent cell motility. Aggregated expression of immunoglobulin genes associated with humoral immunity pathways (*IGHG1, IGHG2, IGHG3, IGHG4, IGKC, IGLC1, IGLC7, IGHM, IGHA1, JCHAIN*) suggests humoral immunity niches of antibody synthesis and secretion in TLS and LNs of pathologic responders (Fig. 5B). Intersecting spots of TLS, islet, and PDAC associated gene expression patterns also showed higher antigen processing and presentation, pointing to local antigen presentation and adaptive immune responses. These observations suggest the development of intratumoral mature TLS is coordinated with localized humoral immunity that could be developed against a mixture of pancreatic islet and tumor antigenic targets.

**Figure 5.**
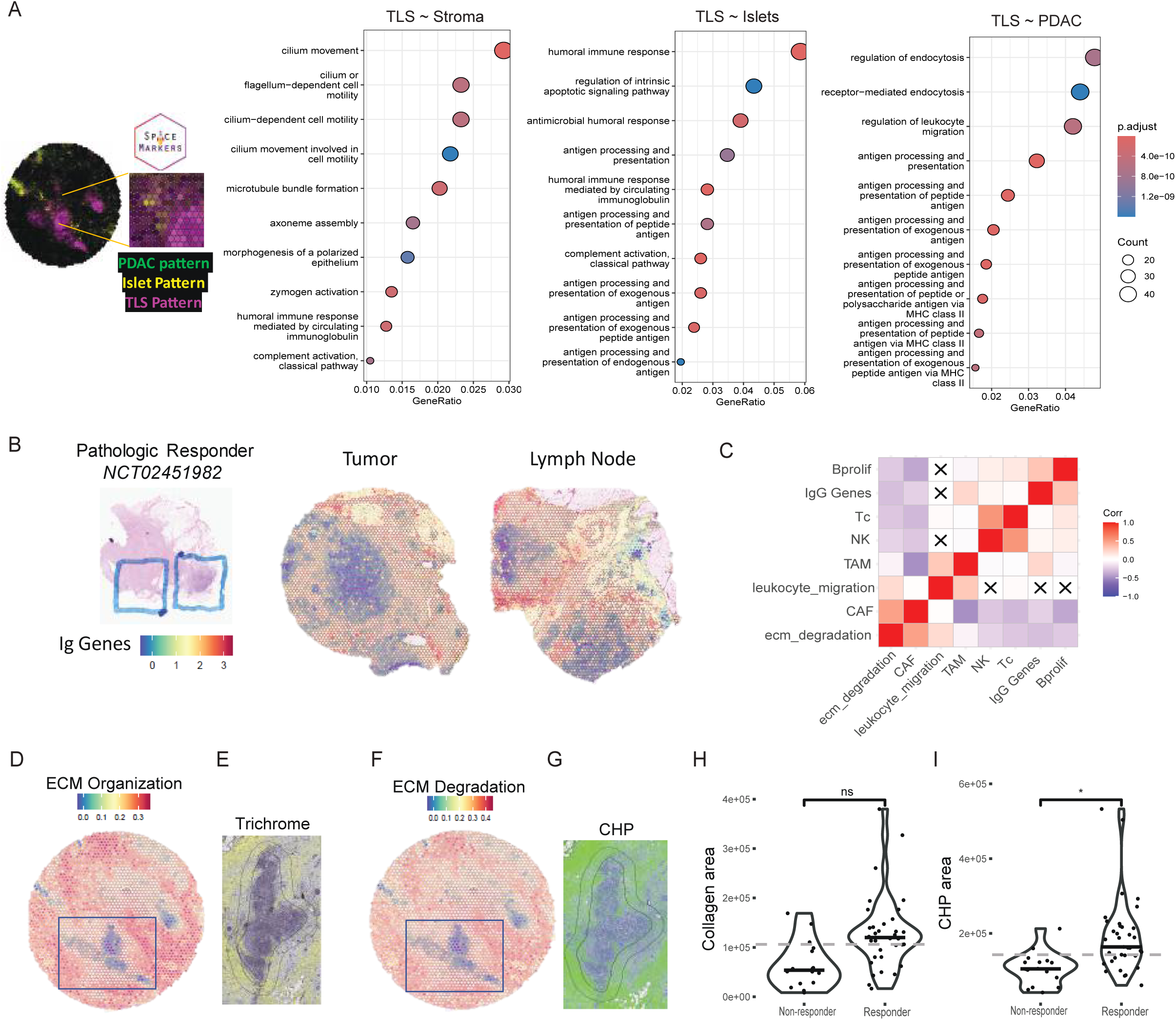
Interactions of spatially variable gene expression patterns demonstrates TLS maturation is coupled with immunoglobulin expression and ECM remodeling. **A**. SpaceMarker analysis of molecular changes from cell-cell interactions showing top enriched pathways at the intersection of TLS and stroma (left), islet (middle), and PDAC (right). **B**. Aggregate expression of Ig genes in a tumor and adjacent lymph node of a pathologic responder. **C**. Correlation plot of IgG genes, leukocyte migration, ECM degradation, and single cell NK, TAM, CAF and B cell projection scores. Crossed out boxes represent insignificant correlations. **D**. Spatial plot showing ECM organization gene set expression in TLS enriched tumor. **E**. Boxed TLS area shown in serial slide stained with trichome and pseudo colored for collagen in yellow. **F**. Spatial plot showing ECM degradation gene set expression in TLS enriched tumor. **G**. Boxed TLS area shown in serial slide stained with collagen hybridizing protein (CHP) and pseudocolored for CHP in green. **H**. Collagen positive area at TLS border in pathologic responders vs. non responders (mixed linear effects testing correcting for individual TLS variation in size, *NS). **I**. CHP positive area at TLS border in pathologic responders vs. non responders (mixed linear effects testing correcting for individual TLS variation in size, *p<0.05).

Based on the SpaceMarkers pathway enrichment results, we hypothesized that plasma cells are activated and trafficking out from mature TLS to produce antibodies that activate the complement cascade and phagocytosis of opsonized cellular targets. The PDAC TME, however, is highly desmoplastic, so we also hypothesized that local immunity would require ECM remodeling to allow immune cell trafficking to targeted tumor areas. We looked at the Pearson correlations of immunoglobulin (Ig) genes, KEGG leukocyte migration, Reactome ECM degradation and organization, as well as single cell reference projections of cytotoxic lymphocytes (CD8 and NK cell), cancer associated fibroblasts (CAFs), tumor associated macrophages (TAMs), and proliferating B cells in ST spots. Proliferating B cells are negatively correlated with ECM degradation and CAFs (Fig 5C). Leukocyte migration is positively correlated with ECM degradation and TAMs (Fig 5C). CAFs are negatively correlated with TAM, NK, Tc, IgG and proliferating B cells, but positively correlated with ECM degradation. IgG gene expression is positively correlated with TAMs and proliferating B cells.

To further investigate the ECM changes and validate remodeling in regions spatially enriched with Reactome ECM organization and degradation gene sets, collagen trichrome staining was performed on adjacent slides (Figs. 5 D-E). This showed pockets of high collagen and low collagen adjacent to the TLS. To determine if these areas of low collagen deposition are actively degraded, we also stained adjacent slides for collagen hybridizing peptide (CHP) (Figs. 5 F-G). Color deconvolution was applied to extract CHP brown and collagen blue positive pixels. Comparison of positive pixels in 200um-wide segments encircling TLS demonstrated similar collagen deposition but higher CHP density in pathologic responders compared to nonresponders (p=0.02) (Figs. 5 H-I). Altogether these results lead us to hypothesize that mature TLS trigger a cascade of events that make the immediately adjacent PDAC TME more permeable to adaptive immune responses.

### Plasma cell spatial dynamics demonstrate trafficking of TLS-associated humoral immunity

Profiling TLS niches in the PDAC TME of pathologic responders uncovered TME remodeling co-occurring with their formation, yet the evolutionary dynamics and targets of their distributed antibody responses are still not clear. Plasma cells are terminally differentiated mature memory B cells that are canonically known to originate from the bone marrow (McHeyzer-Williams and Ahmed, 1999). However, increasing evidence in TLS cancer immunology has demonstrated local plasma cell propagation (Fridman et al., 2023). In our ST data, we observed expression of plasma cell markers, including SDC1 (CD138), in spots surrounding mature TLS (Fig. 3E). The IMC data also showed significantly higher expression of CD138 in germinal center (GC) B cells of TLS regions from pathologic responders compared to non-responders (Supplemental Data S2). We then performed whole-slide multiplex-immunohistochemistry (mIHC) for CD138, PanCK, which stains epithelial cells including cancer cells, CD20 for B cells before they transition into memory plasma cells and CD3 for T cells. We identified CD138^+^ cells localizing between TLS and infiltrating PDAC neoplastic lesions suggesting plasma cell trafficking from the TLS to niche-specific targets (Figs. 6A, 6B).

**Figure 6.**
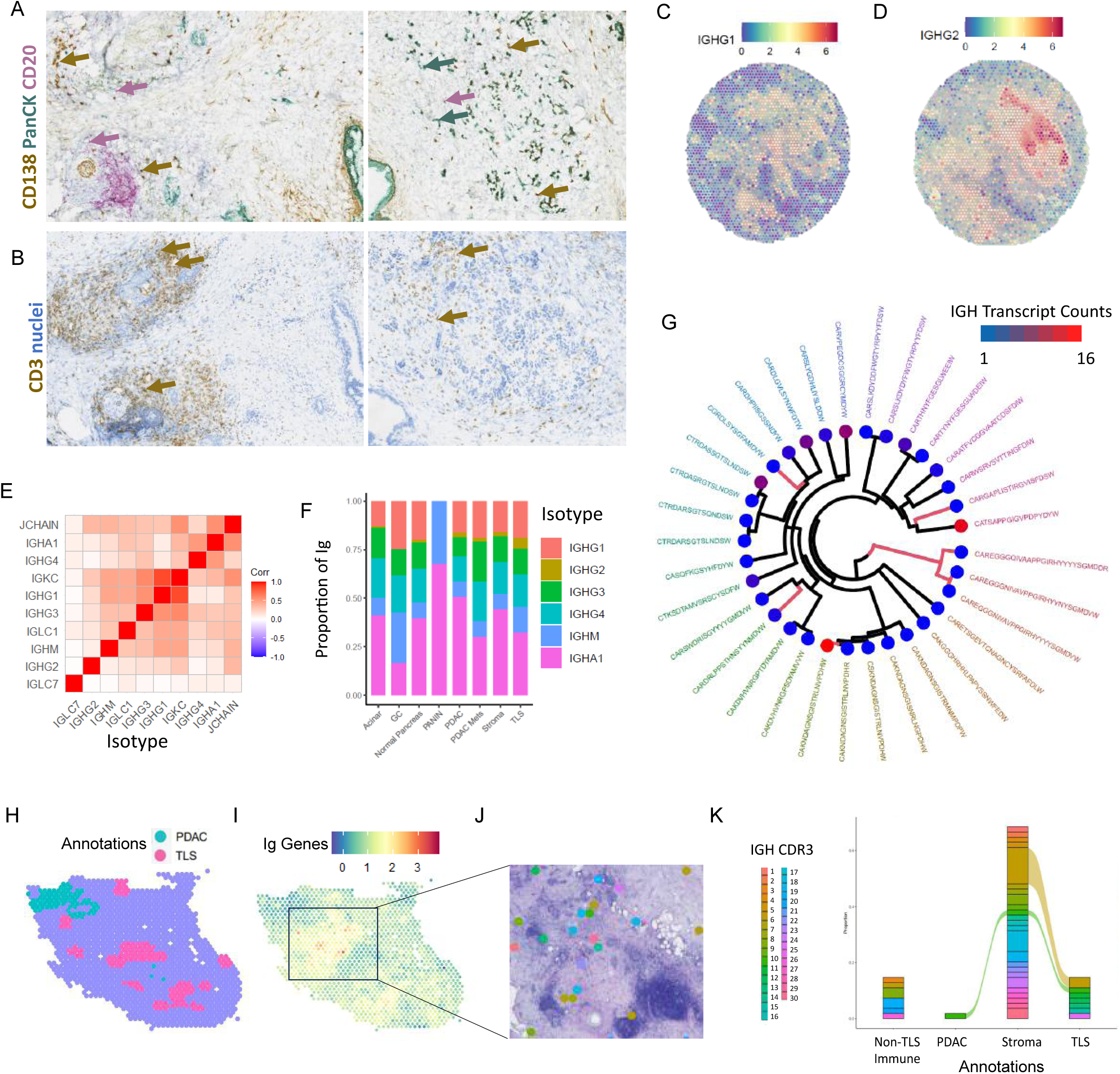
Intratumoral plasma cell dynamics in the TLS-enriched PDAC TME. **A**. Representative triple-stained PDAC samples from this cohort showing infiltration of CD138^+^ plasma cells (brown) in PDAC lesions staining for PanCK (green) and B cells (CD20, magenta). **B**. Serial sections stained for T cells (CD3, brown) and nuclei in counter stain (blue). IGHG1 (**C**) and IGHG2 (**D**) expression in representative sample. **E**. Gene expression correlations of Ig isotype genes. **F**. Stacked barplot showing Ig isotype gene usage by pathology annotations as proportion of total counts of Ig genes expressed per spot. **G**. Dendrogram displaying IGH CDR3 sequences from a representative TLS-dense patient sample (116_1). Position in dendrogram describes hamming distances of CDR3 sequence similarity. Red branches indicate CDR3 sequences found in TLS. Expression of IGH transcript is colored on dendrogram branch endpoints. **H**, Annotations of representative TLS-dense patient tumor sample (116_1) and (**I**) aggregate Ig gene expression overlayed on spatial plot, with top IGH sequences (**J**) plotted in colored dots in zoomed our region with high Ig expression. **K**, Alluvial plot showing representative top IGH chains between PDAC, stroma, and TLS spots.

To further trace these mature B cells, we examined the Pearson correlation of immunoglobulin isotypes in the ST spots, such as IgG1-4, IgG and IgM, highlighting spatial variability among isotypes (Figs. 6C-E). This suggested differential Ig gene isotype usage by site because of Ig class switching in TLS germinal centers. We looked at isotype usage by computing the proportion of each Ig isotype gene expression to total Ig gene expression. Indeed, we identified different isotype usage profiles depending on the nearest histological type (Fig. 6F). Notably, the most prevalent isotype across sites, including TLS, is IgA, except in lymph node germinal centers where the most prevalent is IgM. TLS in this cohort expressed IgG2 which is most commonly considered to be responsive to polysaccharide antigens (Aucouturier et al., 1986).

Isotype switching is coupled with affinity maturation and clonal selection in GCs. The differential Ig gene usage we observed prompted us to investigate B cell clonal dynamics across TLS and associated tissue structures and thus performed B cell receptor (BCR) chain sequence alignment directly from the raw frozen ST sequencing data. Across all samples, 159 Ig Heavy (IGH), 1080 Ig Light (IGL), 1068 Ig Kappa (IGK) unique chains were recovered and clustered in dendrograms using the hamming distance of CDR3 sequences to visualize clonotypes (Fig. 6G, Supplemental Figs. S6A, S6B). Correlating the aggregated clone transcript counts with the transcriptional TLS pattern and IgG score in the ST spots using Pearson’s correlation revealed significantly positive relationships confirming that the aligned BCRs overlap with B cell-related spatial transcriptional programs (Supplemental Figs. S6C). Between patient samples, overlap of IGL CDR3 sequences was observed among the top 50 expanded IGL chains, but not in the top 50 IGH chains (Supplemental Figs. S7A, S7B). Within patient samples, overlap between distinct spatial niches, including TLS, stroma and PDAC was identified, further suggesting TLS-associated local B cell clonal expansion (Figs. 6 H-K, Supplemental Figs. S8A, S8B). Despite the highly desmoplastic and immunosuppressive PDAC TME that is not conducive to lymphocyte trafficking, the immunoglobulin class-switching spatial dynamics shown here in combination with our plasma cell profiling further highlight trafficking of intratumoral B cells in TLS-enriched PDAC.

## DISCUSSION

Our study demonstrates a workflow of *in situ* profiling and computational approaches to study TLS biology using spatial-omics and statistical machine learning. Leveraging these techniques, we navigated the spatial heterogeneity and cellular architecture of the immune landscape in PDAC. Cellular neighborhoods of TLS neogenesis were characterized and their interactions were modelled within the PDAC TME, which demonstrated intratumoral maturation of B cell immunity that is associated with stromal ECM remodeling in pathologic responders.

In a study of PDAC patients receiving chemoimmunotherapy, the presence of a mature TLS gene signature in pretreatment biopsies was shown to be associated with longer survival, while a subset of tumors contained mature TLS that support B and T cell organization through CXCL13 production (Kinker et al., 2023). Although mature TLS rarely form in untreated pancreatic tumors, neoadjuvant immunotherapy in PDAC has shown to induce active intratumoral TLS (Hiraoka et al., 2015; Lutz et al., 2014). In this study, to visualize and deeply characterize TLS in PDAC patients after neoadjuvant immunotherapy, histological evaluations from an expert pancreatic pathologist were incorporated into advanced machine learning techniques. Using the CODA platform which has been previously implemented in 3D reconstruction of PDAC tumors (Kiemen et al., 2022) and histology annotations of spatial transcriptomics of FFPE-embedded pre-malignant pancreatic lesions (Bell et al., 2024), we were able to train a model to quantify various histological classes within the PDAC TME with high accuracy, including putative TLS. Histological classification using machine learning of TLS has been implemented previously in gastric cancers, including three distinct maturation states (Li et al., 2023). In our study, we trained CODA to also learn fiducials in the ST capture areas to facilitate the transfer of histology annotations into ST spots. Subsequently, imaging mass cytometry was implemented to determine TLS maturation in distinct niches. Mature TLS were found in diverse spatial niches, including acinar-rich, islet-rich, stromal, and PDAC cellular neighborhoods. Although the scope of this study was not designed to identify antigen targets, spatial localization of TLS and their immune cells in these niches suggest TLS neogenesis could locally amplify immunity against a range of intratumoral targets that may not be expressed by malignant cells. TLS in islet-rich regions in particular show striking similarities with those found in diabetes, supporting diabetes-like autoimmunity occurring in PDAC tumors after immunotherapy (Korpos et al., 2021; Lee et al., 2006). TLS in acinar niches could be the remnant of an immune response to acinar-to-ductal metaplasia (Kopp et al., 2012), although is not possible to know the exact timing of TLS neogenesis in these patients. An involuted TLS, mature TLS and resolving necrotic duct were co-identified in the same 6.5x6.5mm capture area in a tumor of a pathologic responder. An involuted TLS morphology was first described by Shu et al. (2024) in hepatocellular carcinoma, in which B cell and T cell compartments are flipped in TLS found in tumor resolving regions. This morphology was posed to be part of the end stages of the resolution phase of the TLS lifecycle. Although additional studies are necessary to fully understand the nature and timing of the involuted morphology, our results illustrate parallel TLS lifecycle stages co-occurring during an active TLS-mediated immune response following neoadjuvant treatment, which may be potentially extending the activity span of the response. The diversity of tissue structures in which TLS and associated immune cells infiltrate, as well as the presence of different TLS morphologies in the same tumor region highlight a dynamic immune activation and resolution co-occurring in response to a range of local intratumoral antigens.

To further elucidate the role of TLS in these spatial niches, we applied non-negative matrix factorization to learn sets of co-expressed genes, called spatial patterns, which correlated with pathology annotations. Previous studies have leveraged TLS-associated genes from bulk and single cell data, but not spatial patterns, to build models that demonstrate value in risk assessment and prognostic prediction in PDAC (Hu et al., 2024; Ma et al., 2024). In this study, signatures directly learned from ST data containing TLS projected to TCGA demonstrated their clinical relevance in prognostication of overall survival. The patternMarker genes of the TLS spatial pattern most positively prognostic to survival could be used in subsequent risk-model studies to further estimate the prognostic value of subsets of TLS patternMarkers in different immunotherapy contexts in PDAC and beyond. In this study, we focused on the genes expressed at the intersection of TLS-transcriptional signatures and related spatial patterns in the PDAC TME. SpaceMarkers and ECM degradation analyses demonstrated higher density of humoral immunity activity in TLS niches and ECM remodeling in pathologic responders. The links between humoral immunity, CAFs, stromal remodeling and disease progression in PDAC are not yet well understood. In IPMN samples, spatial transcriptomics and proteomics have shown an incremental decrease of plasma cell infiltration as dysplasia and cancer progress (Sans et al., 2023). Studies of CAFs in PDAC have shown the role of distinct CAF subsets in stromal remodeling, such as α-SMA^+^ and FAP^+^ CAFs in in well and poorly differentiated cancer niches, respectively (Sherman and Beatty, 2023). Although beyond the scope of this paper, mechanistic exploration of TLS-derived humoral immunity and CAF crosstalk could present a new avenue for therapeutic exploration in ameliorating the desmoplastic PDAC TME.

In several cancers including NSCLC, melanoma, and sarcoma renal cell carcinoma, plasma cell infiltration has been associated with greater survival and improved responses to PD1/PDL1 immunotherapy (Teillaud et al., 2024). Germinal center reactions in TLS and IgG have also been associated with improved survivorship in treatment-naïve PDAC using bulk RNA-seq and multiplex IHC (Gunderson et al., 2021). In our study treating PDAC patients with combination neoadjuvant immunotherapy (GVAX, aPD1, a41BB), the spatial distribution of isotype gene usage and BCR clonotypes in the ST data inferred class switching and plasma cell trafficking from TLS into the PDAC TME. Using plasma cell markers we confirmed their infiltration in PDAC niches. Although hotspots of high IgA and IgG gene expression in TLS niches were observed, the epitopes targeted by locally expressed IgA and IgG antibodies in pathological responders in PDAC with intratumoral TLS remain to be explored. In a study deploying IgG sequencing, antibody reconstruction and antigen discovery in PDAC found plasma cells secreting autoantibodies, notably F-actin), the nucleic RUVBL2, and heat shock proteins Hsp60/HSPD1 (Yao et al., 2023). However, the TLS sequenced in this study were exclusively found in the tumor border, while IgA targets were not explored. In another study, tumor cell-derived debris and IgG were found to promote disease progression in PDAC TAM-induced inflammation (Chen et al. 2019). In renal cell carcinoma, tumors with TLS and IgG-bound tumor cells were found to be infiltrated by plasma cells producing IgG and IgA along fibroblastic tracks, proposing complement (CDC) or antibody-dependent cell cytotoxicity (ADCC) (Meylan et al., 2022). In our study, we indeed also found enriched complement activation associated with TLS using SpaceMarkers as well as NK-cell signature colocalized with TLS. However, we cannot experimentally determine a CDC or ADCC mechanisms in this body of work, as additional NK and TAM profiling markers in IMC or equivalent would be necessary to elucidate such processes *in situ*. Overall, our results contribute to growing evidence that combinations of neoadjuvant immunotherapies enhance the formation of mature TLS and highlight the role of intratumoral B cells in propagating local antigen production and coordinating TME reprogramming.

Our study has some technological and practical limitations. The spatial multi-omics datasets from neoadjuvant immunotherapies presented here represent a two-dimensional static snapshot of TLS development in patient samples at a specific time point after treatment. Novel experimental mouse models that can fully recapitulate human TLS biology in PDAC could serve as a future platform to study temporal dynamics, while future comprehensive 3D profiling technologies could expand current 2D dimensional models. Additionally, although the scope, design and technologies used in this study were not intended for antigen discovery, much of the analysis of intratumoral B cell trafficking performed here is limited to spatial correlations. Future reconstitution of intratumoral BCR and antibody-antigen target discovery studies would offer more compelling evidence regarding the nature of the humoral immunity in the PDAC tumors studied here.

Our spatial investigation of TLS within resected pancreatic tumors from patients treated with neoadjuvant immunotherapy revealed distinct cellular neighborhoods which were integral to understanding the spatial dynamics of TLS in the PDAC TME. The analysis described in this study focused on spatial immune dynamics of intratumoral TLS in PDAC and provides unprecedented spatial molecular dimensionality of the PDAC tumor immune microenvironment. The transcriptional spatial patterns we learned could be used in future targeted panel design that will resolve the TME at single cell or subcellular resolution, while the 28-sample spatial transcriptomic dataset generated here could be leveraged in future studies examining other aspects of PDAC and pancreas-associated lymph node pathology. Based on the results of our analysis, we propose that next-generation TLS induction immunotherapy strategies should be designed with the intent to induce a niche-bias that would further augment pathologic responses into long-term clinical responses. A deeper understanding of intratumoral B cells and antibodies could strengthen our understanding of the role of autoantibodies in PDAC and lead to novel therapeutic approaches.

## METHODS

### Fresh frozen cohort spatial transcriptomics data generation and preprocessing

For the frozen cohort, a total of 16 pancreatic cancer samples resected from GVAX-treated PDAC patients (clinical trial #NCT00727441) were embedded in OCT media and freshly frozen. Sections were cut into 6.5 x 6.5 mm dimensions compatible with Visium spatial transcriptomics slides. The frozen cohort contained 14 tumor sections and 2 adjacent lymph nodes. We used 4 Visium slides and covered the 16 capture areas with sections cut in a cryostat at 10 µm. The fresh frozen Visium protocol was followed according to manufacturer’s recommendations (10x Genomics Spatial 3’ v1). Tissue permeabilization and library construction was carried out according to the manufacturer’s protocols. Permeabilization time was optimized to 30 minutes. The captured tissue sections were permeabilized to allow RNA to be captured onto the slide where it reverse-transcribed to cDNA. The cDNA was then amplified, and sequencing libraries were prepared using the 10x Genomics Visium Spatial Gene Expression Reagent Kit for Library Preparation. The libraries were then sequenced using an Illumina NovaSeq sequencer. The raw data obtained from the sequencing were processed using the SpaceRanger (spaceranger-1.2.0) pipeline for alignment onto the GRCh38-2020-A reference transcriptome quantification of gene expression. The resulting aggregated expression matrices were imported into Seurat (version 4.0.1) for downstream analysis in R version v4.0.2. The data were normalized using SCT tranform function in Seurat to account for differences in library size and to minimize the impact of technical noise. Certified pathology annotations were manually performed and incorporated in the 10x Loupe browser.

### FFPE cohort spatial transcriptomics data generation and preprocessing

For the FFPE cohort, a total of 12 FFPE archival samples from a neoadjuvant immunotherapy combination trial in PDAC (clinical trial #NCT02451982, GVAX n=5, GVAX+aPD1 n=2, GVAX+aPD1+a41BB n=5) were used for area selection for sequencing. All samples selected were checked for RNA quality checked by isolating 20um sections of each sample using the RNase FFPE kit (Qiagen), following manufacturer’s instructions. RNA quality was measured using the DV200 assay on the Bioanalyzer (Agilent) to determine the proportion of fragments with ∼200bp in the sample. RNA quality was considered good if DV200 > 50%. After sectioning, expert pathologist selected 8 high-quality 6.5x6.5 mm representative regions of pathologic response regions in tumors and 4 regions in associated tumor-adjacent lymph nodes. Using the Visium FFPE (10x Genomics) platform and following manufacturer’s validated protocol the samples were deparaffinized, stained with hematoxylin (discovery cohort) or H&E (validation cohort), and scanned using the Nanozoomer scanner (Hamamatsu) at 40x magnification. Human probe hybridization was performed overnight at 50°C. Following probe ligation, the RNA was digested, and the tissue was permeabilized for the release, capture, and extension of the probes. The designated area for each sample is covered by spatial capture probes containing oligo-d(T) that hybridize to the poly-A tail sequence present in the transcript-targeting capture probes. The sequencing library preparations were performed as instructed by the manufacturer using the extended probes as the template. Libraries were sequenced with a depth of at least 50,000 reads per spot (minimum of ∼250 million/sample) at the NovaSeq (Illumina). The Visium Human Transcriptome Probe Set v1.0 contains probes to 19,144 genes and after computational preprocessing (filtering for probes off-target activity) provides gene expression information for 17,943 genes. The raw sequencing data were processed using the SpaceRanger Space Ranger was v2.1.0 pipeline for alignment onto the GRCh38-2020-A reference transcriptome quantification of gene expression. The resulting aggregated expression matrices were imported into Seurat (v4.0.1) for downstream analysis in R v4.0.2. The data were normalized using SCT tranform function in Seurat to account for differences in library size and to minimize the impact of technical noise.

### CODA H&E classification model training and integration with FFPE ST

A machine learning model for H&E image segmentation for PDAC H&E sections on 10x Visium ST slides following the CODA implementation strategy described in Bell et al., (2024). Briefly, eleven microanatomical components of human pancreas tissue were multi-labelled using a semantic segmentation workflow as described by Kiemen et al. (2022). The components recognized were: 1) islets of Langerhans, 2) PDAC, 3) normal ductal epithelium, 4) ECM, (4) fat, 5) Visium fiducials, 6) TLS, 7) acini, 8) nerves, 9) fat, 10) whitespace, 11) vasculature. A minimum of 50 manual annotations per slide with representation from each tissue type (where available) were labelled on H&E and corroborated with expert pathologist using Aperio ImageScope. In each slide, half of the newly generated annotations were used in the training dataset for the convolutional neural network and the other half were used as an independent testing dataset to evaluate model performance. Following training, the tissue images were segmented at a resolution of 1µm per pixel. Nuclear coordinates were generated via the detection of two-dimensional intensity peaks in the hematoxylin channel of the deconvolved H&E images. Briefly, the TMA images were down sampled to 1 µm/pixel resolution. To adapt CODA to the hematoxylin only stained images (test cohort), the color image was converted to greyscale. No changes were necessary for the H&E-stained sections (validation cohort). The image was smoothed using a Gaussian filter and two-dimensional intensity peaks with minimum radii of 2µm were identified as nuclear coordinates. The low-resolution image used for the Visium pre-processing with Space Ranger was registered to the high-resolution tissue image used for microanatomical measurements to integrate the two workflows. The registration utilized the fiducial markers present on the ST glass slide to estimate the registration scale factor and translation. As registration was performed on two scans of identical tissue sections, it was assumed that rotation was not necessary. Here, the low-resolution image was registered to the high-resolution image (rather than the other way round) so that the scale factor was always greater than 1 and ensuring that the 1 µm resolution of the tissue micro annotations was preserved. First, the fiducial markers in each pair of images were segmented by identification of small, nonwhite objects surrounding the larger TMAs. Nonwhite objects were determined to be pixels with red-green-blue standard deviations greater than 6 in 8-bit space. These objects were morphologically closed, and very small noise (<50pixels) were removed. The fiducial markers were then determined to be objects in the image within 20% of the median object size (as many fiducial markers existed for each corresponding tissue image). This process resulted in fiducial image masks for the high-resolution and low-resolution tissue images. With these masks, four possible registrations were calculated to account for the situation where the Visium analysis was performed on the tissue image rotated at a 0-, 90-, 180-, or 270-degree angle. For each registration, the corner fiducial markers of the low-resolution image were rescaled and translated to minimize the Euclidean distance to the fiducial markers of the high-resolution image. Of the four registration results, the registration resulting in the greatest Jaccard coefficient between the high-resolution and low-resolution fiducial masks was chosen. For the twelve TMAs, the average Jaccard coefficient of the fiducial masks was 0.94. The registration information used to overlay the low-resolution tissue image to the high-resolution tissue image was used to convert the coordinates corresponding to the location of the Visium assessment in the low-resolution image into the high-resolution images coordinate system. Once the Visium coordinates were registered to the high-resolution image, the generated tissue microanatomy composition and cellularity were calculated for regions within 25µm of each coordinate. For each Visium coordinate, pixels in the micro-anatomically labelled mask image within 25µm of that coordinate were extracted. Tissue composition was determined by analyzing the % of each classified tissue type within that dot. The cellularity of each dot was determined by counting the number of nuclear coordinates within 25µm of each Visium coordinate. Cellular identity was estimated by determining the microanatomical label at each coordinate where a nucleus was detected. For ST spot annotations, the class with the highest proportion of classified pixels in each spot was used for labelling.

### Nonnegative matrix factorization of ST data and spatial patterns analysis

The CoGAPS R package (v3.58) was used for NMF pattern learning with the following parameters: distributed = “genome-wide”, sparseOptimization = TRUE, and defaults for the remaining parameters (Fertig et al., 2010). CoGAPS runs were set up with an increasing number of patterns. For spatial analyses, CoGAPS objects with 10 and 15 patterns were chosen, for the frozen and FFPE ST data respectively. Spatial patterns were projected into a PDAC single cell reference Atlas (Guinn et al., 2024) and The Cancer Genome Atlas (TCGA) bulk RNAseq data using the projectR R package v1.20.0 (Stein-O’Brien et al., 2019). Survival analyses were performed across all TCGA cancer subtypes and in pancreatic cancer separately (PAAD). In PAAD, we used a previously curated version of the cohort which accounted for normal samples and neuroendocrine tumors (Peran et al., 2018). Clinical and gene expression data from the TCGA database were downloaded in R from the Genomic Data Commons (GDC) Data Portal using the TCGAbiolinks R package v2.32.0 (Colaprico et al., 2016). The clinical data included overall survival (OS) time, censoring status, and relevant demographic and clinical covariates. Gene expression data were normalized using the fragments Per Kilobase of transcript per Million mapped reads (FPKM) method and log2-transformed for downstream analyses. Kaplan-Meier (KM) survival curves were generated to estimate the survival probabilities across different patient subgroups based on the level of projected ST pattern enrichment in bulk RNAseq data. Patients were stratified into high and low expression groups based on the top 5^th^ percentile expression value of the pattern of interest. Survival curves were visualized using the survminer R package (v0.4.9), and log-rank test p-values < 0.05 were considered statistically significant. Univariate Cox proportional hazards regression was performed to evaluate the association between ST pattern projection weights and overall survival. The Cox model was fitted using the survival R package (v3.5), with hazard ratios (HR) and 95% confidence intervals (CI) reported for each covariate. ST spots spatially enriched for each pattern (hotspots) and differential expression analysis between patterns of interest were computed using the SpaceMarkers R package (v1.0) (Deshpande et al., 2023). SpaceMarkers differential expression between interacting regions and non-interacting regions for each pattern pair combination was implemented on each ST sample in SpInMarkersMode = “residual” mode using CoGAPS patterns as input. Differentially expressed genes were interpreted using the enrichGO and enrichKEGG functions of clusterprofiler version 4.12.6 using default parameters. Correlation plots were generated using the corrplot package (version 0.92), with Pearson’s correlation coefficient employed to assess the linear relationships between Patterns and other continuous variables.

### Spatial immune repertoire analysis

MiXCR (v3.0.12, 20) (Bolotin et al., 2015) was used to align BCR sequences in raw gene expression sequencing FASTQ files from the ST data (frozen cohort only due to compatilble 3’ mRNA capture chemistry). The MiXCR analyze shotgun function with parameters -s hsa \ --starting-material rna \ --only-productive \ --impute-germline-on-export \-- contig-assembly \ --assemble “-ObadQualityThreshold=0.” Aligned chains from IGH, IGK and IGL loci were incorporated into the ST Seurat object and visualized with Seurat. Dendrograms were generated using package ape v5.8. Spatial Alluvial plots of IGH chains showing overlap between annotated ST spots were visualized using ggalluvial extension of ggplot2 (version 3.3.5). Correlation plots were generated using the corrplot package (version 0.92), with Pearson’s correlation coefficient employed to assess the linear relationships between Isotypes.

### Imaging Mass Cytometry (IMC) data acquisition and analysis in FFPE cohort

Serial adjacent slides to the ST FFPE cohort were used for IMC imaging. Staining was done as previously described by Ho et al (2021). Briefly, formalin-fixed paraffin-embedded (FFPE) resected PDAC tissue sections were baked and deparaffinized in histological-grade xylene. Slides were rehydrated with a descending ethanol gradient (100% EtOH, 95% EtOH, 80% EtOH, 70% EtOH) in Maxpar Water (Standard BioTools). Slides were incubated with Antigen Retrieval Agent pH 9 (Agilent® PN S2367) at 96 °C for 1 hour then blocked with 3% BSA for 45 min at room temperature followed by overnight staining at 4°C with the antibody cocktail (see Supplemental Table S1 for list of antibody-metal conjugates). Images were acquired using a Hyperion Imaging System (Standard BioTools) at the Johns Hopkins Mass Cytometry Facility. A total of 23 tiles of ROIs of minimum 1 mm^2^ were selected for imaging from stained slides. Upon image acquisition, representative images were visualized and generated through MCD™ Viewer (Standard BioTools). Images were segmented into a single-cell dataset while retaining spatial coordinates and registering all nearest neighbors for each individual cell using the publicly available software pipeline based on CellProfiler, ilastik, and HistoCAT (Berg et al., 2019; Carpenter et al., 2006; Schapiro et al., 2017). Single cells were clustered using FlowSOM (Van Gassen et al., 2015) into 15 metaclusters, which were collapsed into 10 clusters based on similar gene expression profiles and annotated into final cell types using canonical marker expression. Differential expression analysis of markers in each cell types between groups was performed with Wilcoxon tests and FDR-adjusted pvalues were considered significant at a threshold of 0.05 or below. Composition analyses were normalized to total cell count per group for each cell cluster. All statistical analyses were performed in R version v4.0.2. Spatial plots, boxplots, composition barplots and violin plots were visualized using ggplot2 (version 3.3.5) plotting functions. IMC metadata for each tile ablated are provided in Supplemental Data S1.

### Immunohistochemistry on frozen samples

Automated triple staining was performed on the Leica Bond RX (Leica Biosystems, Buffalo Grove, IL). Heat induced antigen retrieval was not performed in the first round. Endogenous peroxidase was blocked using Peroxidase block (Refine Kit) then anti-CD138 clone MI15 (M7228, Agilent Technologies Inc., Santa Clara, CA) was applied for 30 min at a concentration of 0.108 µg/mL at room temperature using Antibody Diluent (S302283-2, Agilent Technologies Inc). Detection was performed using the Bond Polymer Refine Kit (DS9800, Leica Biosystems). A round of low pH sodium citrate buffer antigen retrieval for 20 min at 95°C was performed. Anti-CD20 clone L26 (CD20-L26-L-CE, Leica Biosystems) for 15 min at a concentration of 0.19 µg/mL using Antibody Diluent. Detection was performed using the Bond Polymer Red Refine Kit (DS9390, Leica Biosystems). A second round of low pH sodium citrate buffer antigen retrieval for 20 min at 95°C was performed. Anti-panCK clone AE1/AE3 (M3515, Agilent Technologies Inc) for 15 min at a concentration of 0.654 µg/mL using Antibody Diluent. Detection was performed using the Green Chromogen (DC9913, Leica Biosystems). Slides were counterstained, baked and coverslipped using Ecomount (5082832, Biocare Medical, Walnut Creek, CA).

### Collagen remodeling analysis on FFPE cohort

To visualize collagen content in the specimens examined by ST, trichrome staining was performed according to Harper et al., (Harper et al., 2022). Briefly, sections were baked at 65°C for 1 hr and de-paraffinized by xylene then rehydrated and stained with a trichrome stain kit (Abcam). Briefly, slides were incubated in Bouin’s fluid overnight, washed with water, then stained with Weigert’s Iron Hematoxylin for 12 minutes. Slides were washed with water then stained with Biebrich Scarlett/Acid Fuschin solution for 10 minutes, then washed with water. Slides were then differentiated in phosphomolybdic/phosphotungstic acid for 15 minutes, changing to fresh solution halfway through. Lastly, slides were stained with aniline blue for 15 minutes, washed with water, differentiated in 1% acetic acid for 10 minutes before dehydration in ascending concentrations of ethanol followed by xylene, and slides were mounted with Cytoseal XYL. To visualize and quantify areas of degraded collagen on the same specimens, collagen hybridizing peptide staining was performed according to Harper et al. 2022. Briefly, sections were de-paraffinized by xylene and rehydrated in decreasing amounts of ethanol. Endogenous peroxidases were blocked with 3% H2O2. Slides were blocked with avidin/biotin blocking kit (Vector Laboratories, SP-2001), then blocked for 1 hour with 3% BSA in phosphate buffered solution (PBS). The collagen hybridizing peptide biotin conjugate (B-CHP, 3Helix, BIO300) was heated at 80°C for 5 minutes, cooled on ice for 30 seconds, diluted to 0.5μM in PBS, added to slides and incubated at 4°C overnight. Slides were washed with PBS and treated with Vectastain ABC kit (HRP) (Vector Laboratories) according to manufacturer specifications, developed with 3,3-diaminobenzene and counterstained with hematoxylin. Tissues were dehydrated in increasing concentrations of ethanol followed by xylene and mounted with Cytoseal XYL. Slides were scanned using Hamamatsu slide scanner and annotated with ImageScope software. TLSs were identified using CODA on serial H&E slides and subsequently annotated on the trichrome and CHP images manually. Collagen and CHP areas were calculated from the respective images by extending these annotations to a radius of 200μm and calculating the positive pixel area for the stains of interest. Collagen and CHP percentages were calculated as the percent of positive pixels within the 200μm radius of the boundary of the annotation. We assumed that the collagen and CHP content surrounding TLSs would differ by pathological response and assumed contributions of random effects by patient. Therefore, we fit a linear mixed-effects model using the ‘lme4’ R package. Analysis of Variance (ANOVA) was employed to compare the fixed effect of the pathological response on the collagen and CHP content. P-values were computed using the ‘lmerTest’ R package. The analysis was conducted in R version 4.4.0, with ‘lme4’ version 1.1-35.3, and ‘lmerTest’ version 3.1-3.

## Data and code Availability

Data analysis code available under github repository github.com//Dimitri-Sid/PDAC_TLS. Raw sequencing data and clinical metadata can be found at dbGaP under controlled access. Processed transcriptional counts data from spaceranger are shared on GEO under submission GSE277116. IMC proteomics data are shared on Zenodo under doi 10.5281/zenodo.13764100. All sequencing data and associated clinical data will be made available after a formal request to the NIH Data Access Committee. dbGaP, GEO and Zenodo portals will be made available upon publication.

## Ethics and institutional review board

Experiments on human samples in this study were performed under the explicit written consent of the patients enrolled under clinical trial trials NCT00727441 and NCT02451982 and the approval of the institutional review board at Johns Hopkins Medicine.

## Author Contributions

DNS, EJF, LTK, RA, LZ, EMJ, conceived of the project. DNS, LTK, RA, QZ designed and conducted spatial transcriptomics experiments. XY, WJH, SS conducted IMC experiments. Data curation and computational analyses were conducted by DNS, AG, MW. ECM stains were conducted by EIH and analysis by DB, ATW and DNS. IHC stains on frozen samples were performed by VJ and AO. CODA model was trained and implemented by DNS, LDe, AK. RA was the expert GI pathologist, reviewed all specimens and provided pathology annotations. DNS, EJF, LTK drafted original manuscript. Clinical samples were provided by LZ. Funding was acquired by EJF, LTK, EMJ. LDa conducted code review. DNS and DL deposited data. All authors reviewed manuscript.

## Acknowledgments

The authors would like to thank: the patients and their families; the Oncology Tissue Services (OTS) Core Facility and the Reference Histology Lab at JHU for tissue sectioning services; the Genetic Resources Core Facility (GRCF) library sequencing services. Graphical abstract was made in biorender.com.

This work was supported by the following sponsors: The Lustgarten Foundation Pancreatic Cancer Research grant (to E.M.J., E.J.F and L.T.K), The Sol Goldman Pancreatic Cancer Research Center grant (to L.T.K.), NIH P01-CA247886-01A1 (to E.M.J., E.J.F and L.T.K), SU2C/AACR DT-14-14 (to E.M.J.), the Emerson Cancer Research Fund (to E.M.J., E.J.F.), an Allegheny Health Network (AHN) grant (to E.J.F.), NIH U01CA212007 (to E.J.F.), NIH U01CA253403 (to E.J.F.), NCI R50 CA243627 (to L.Da) the JHU Discovery Award (to E.J.F.), and SPORE GI P50CA062924-24A1 (to E.M.J, E.J.F. and L.T.K), NCI F31CA268724-01 (to D.N.S), Sanofi (W.J.H.), NeoTX (W.J.H.), E.I.H. is supported by K00AG068527 (NIA). A.T.W. is supported by a Team Science Award from the Melanoma Research Alliance, Samuel Waxman Research Foundation and the Mark Foundation. A.T.W. is also supported by P01CA114046 (NCI), a Bloomberg Distinguished Professorship and the EV McCollum Endowed Chair.

