## Supplemental Figures and Tables for "Spatial multi-omics reveal intratumoral humoral immunity niches associated with tertiary lymphoid structures in pancreatic cancer immunotherapy pathologic responders"

**Supplemental Figure S1.** Spatial correlations of markers in intratumoral TLS of responders (**A**) and nonresponders (**B**). **C**, Heatmap illustrating the 10 single cell clusters and their expression of protein markers as well as their overall abundance in the IMC dataset.

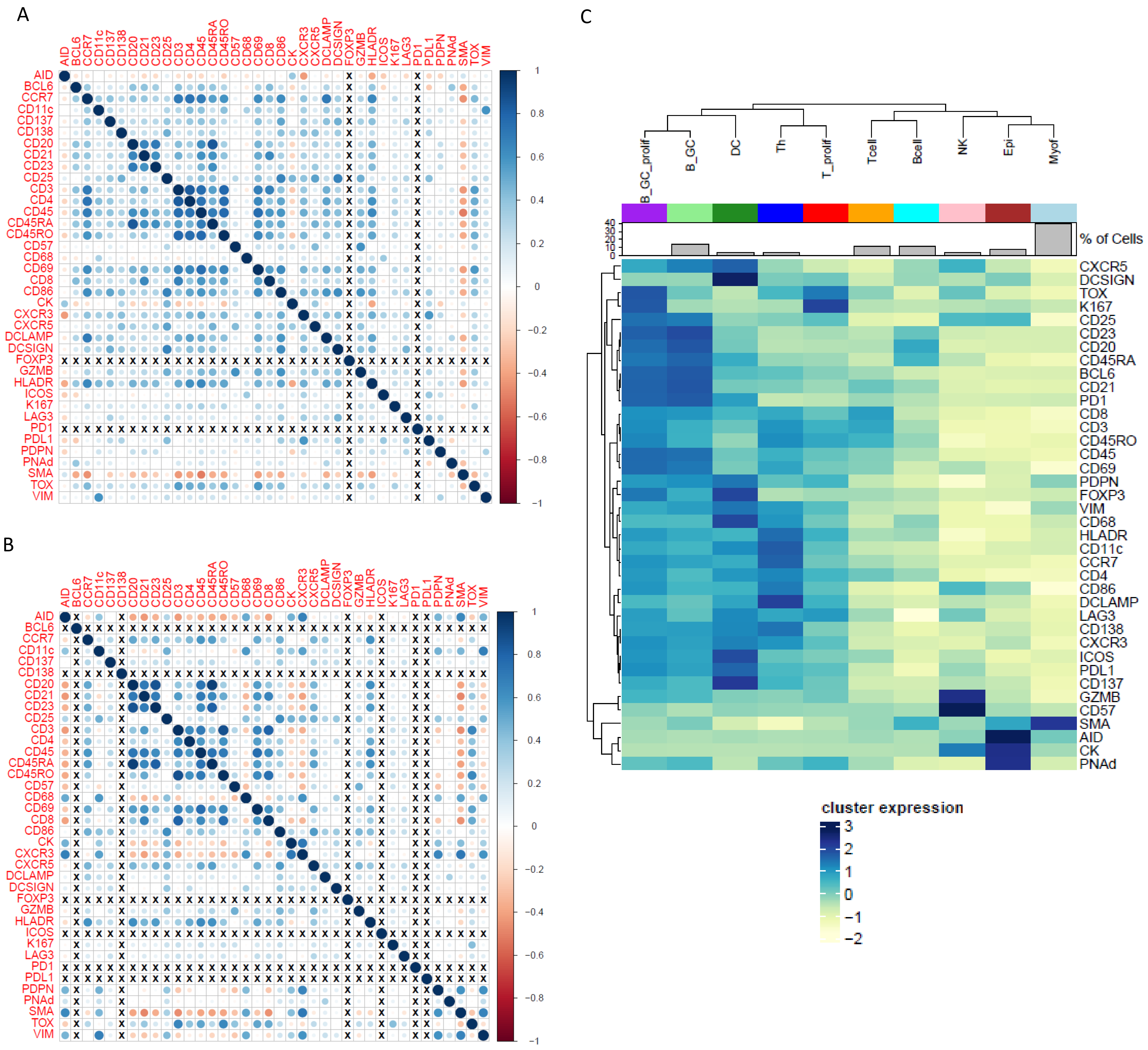

**Supplemental Figure S2.** Imaging mass cytometry single cell segmentation results of intratumoral TLS, peritumoral TLS and tumor adjacent lymph nodes (LN). **A**, Spatial plots showing spatial distribution of single cells after clustering. **B**, Stacked barplot of cell type composition by IMC tile. **C**, Stacked barplot showing aggregated compositions comparing tiles from responders to non-responders in each sample type.

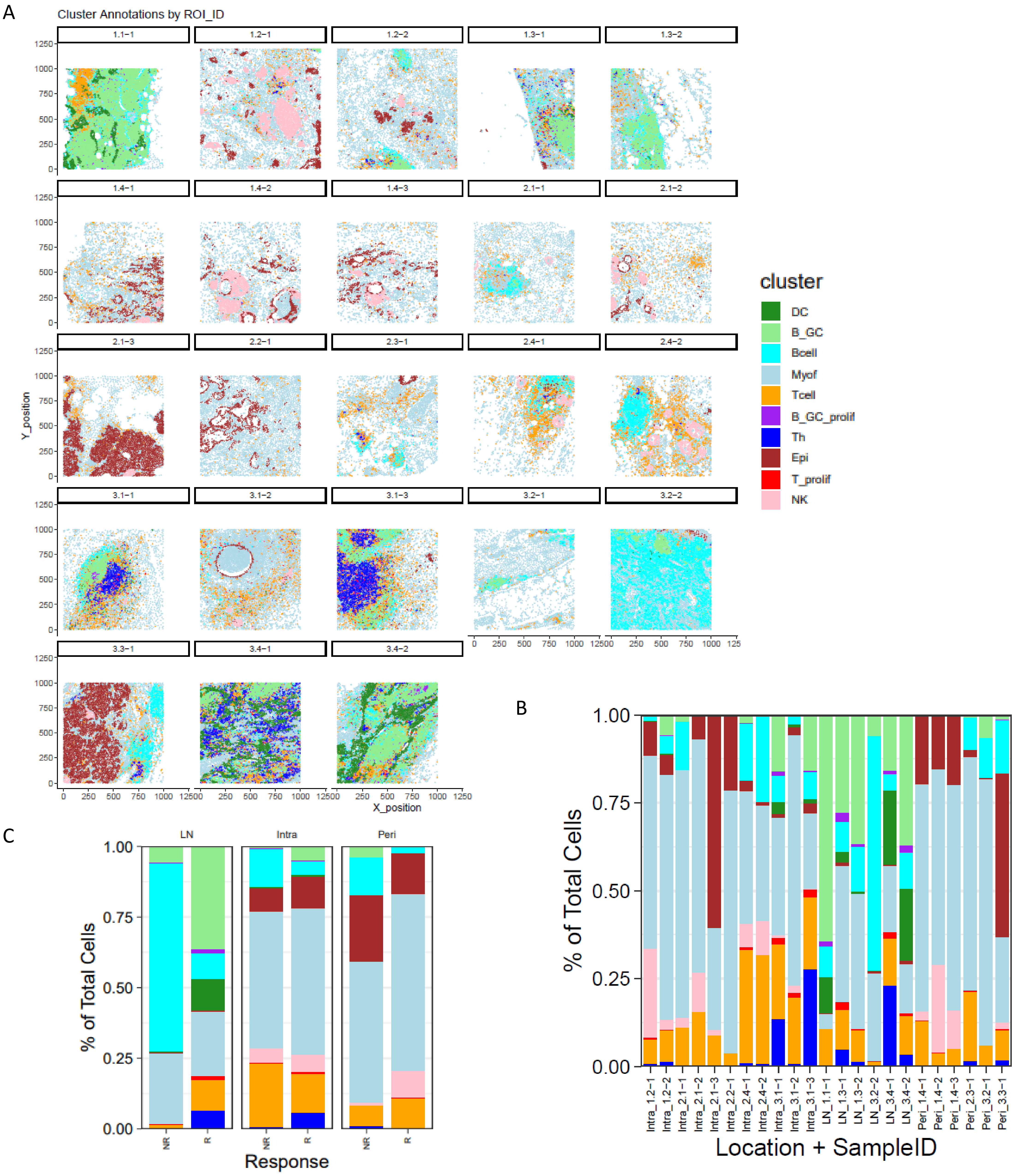

**Supplemental Figure S3.** Intratumoral TLS neighborhood analysis in responders (**A**) and nonresponders (**B**). Composition pie charts showing the top neighbors by cell type.

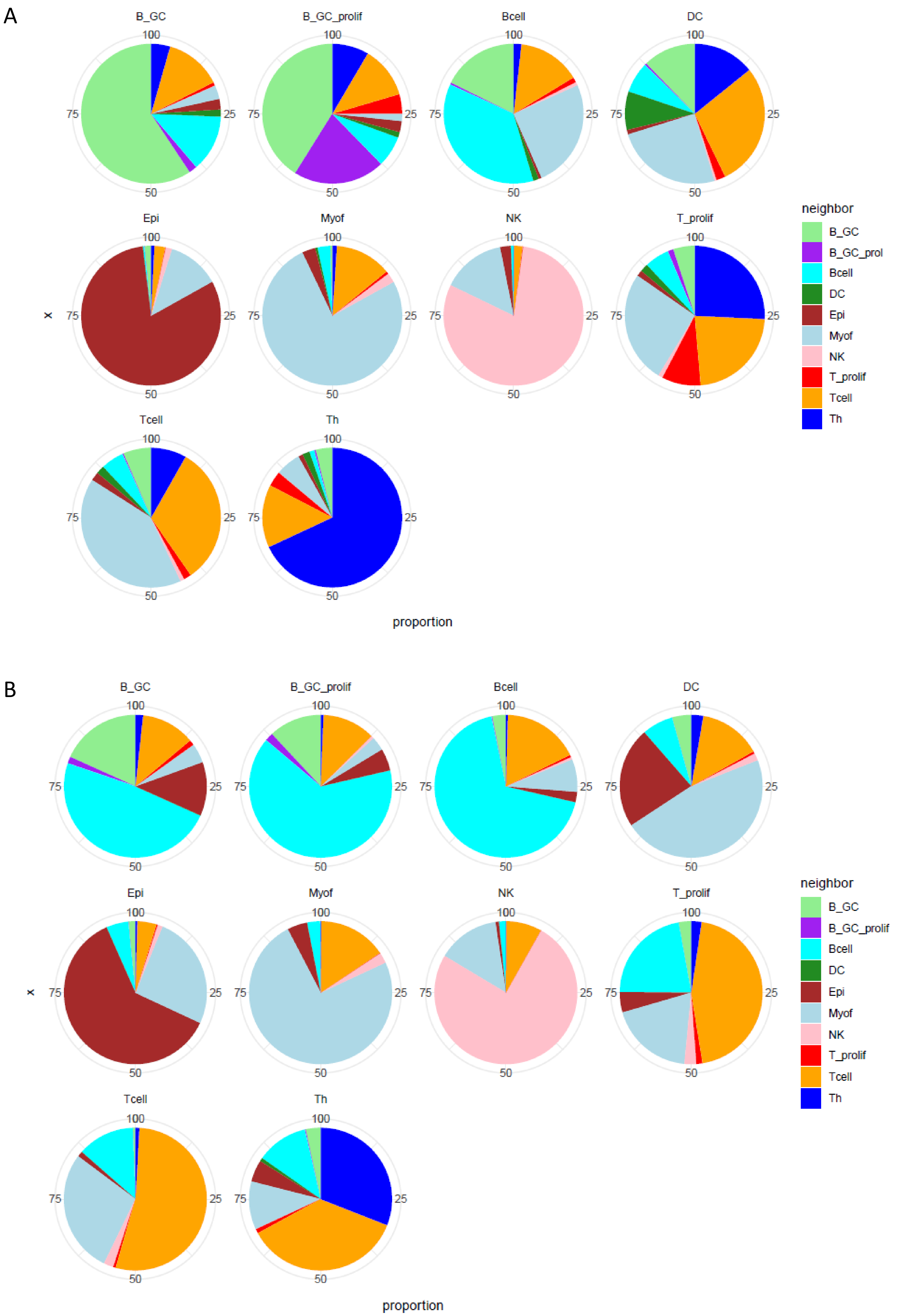

**Supplemental Figure S4.** Transferring histological annotations and nuclei registration from CODA model trained on PDAC TME to Visium spatial transcriptomics spots. Spatial plots showing the percent of TLS-annotated pixels (**A**) and nuclei (**B**) in spots overlayed with histology in FFPE cohort. **C**, top annotation per spots as computed by annotated pixels and nuclei. **D**, total registered nuclei by annotation class.

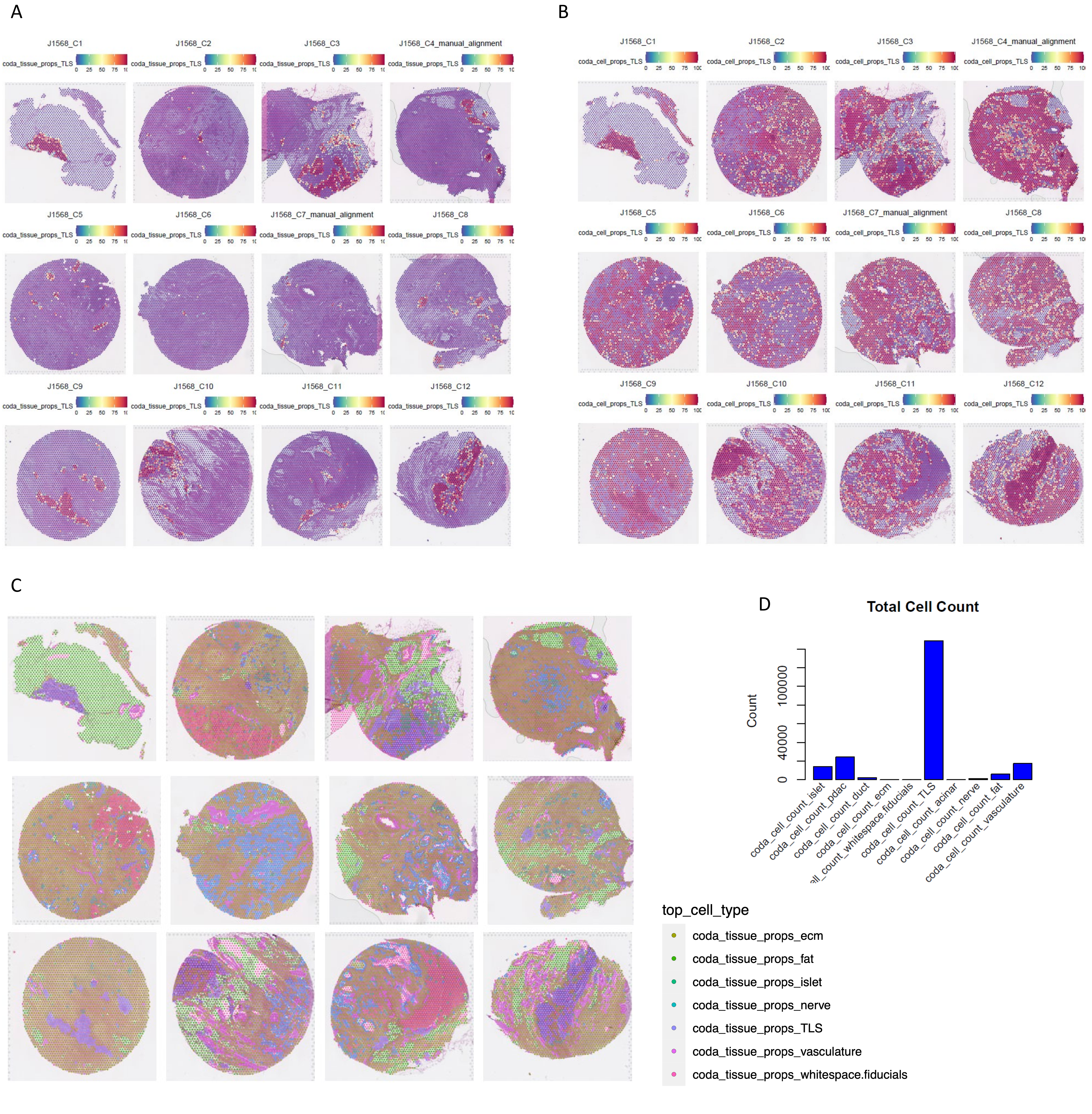

**Supplemental Figure S5.** Forrest plot of cox regression model of Frozen ST spatial patterns on TCGA pancancer bulk RNAseq data.

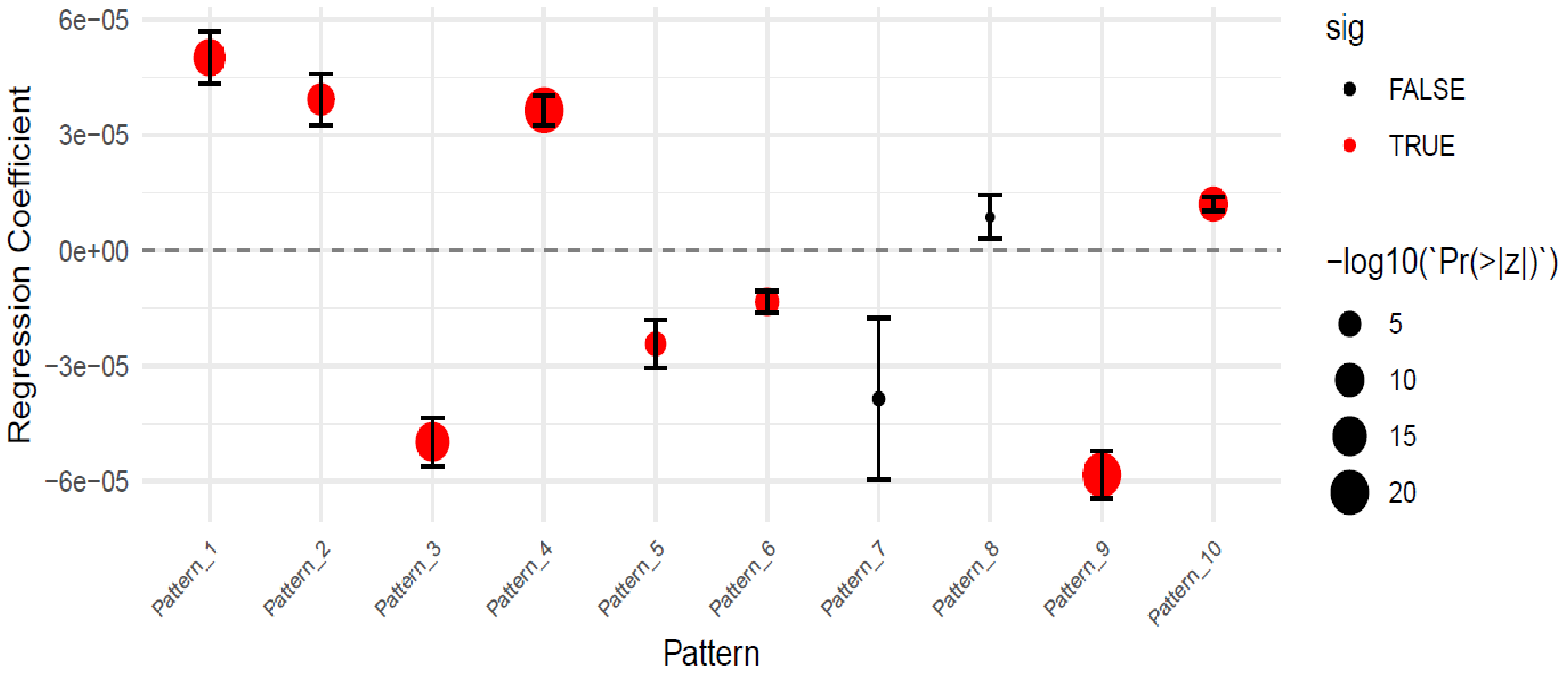

**Supplemental Figure S6.** Dendrograms of IGH chains across dataset coloring branches PDAC (**A**) or TLS (**B**) annotations in red. **C**, Spatial correlations of NMF patterns, aggregated IgG gene expression (iggscore) and total BCR counts (CloneCoutSums).

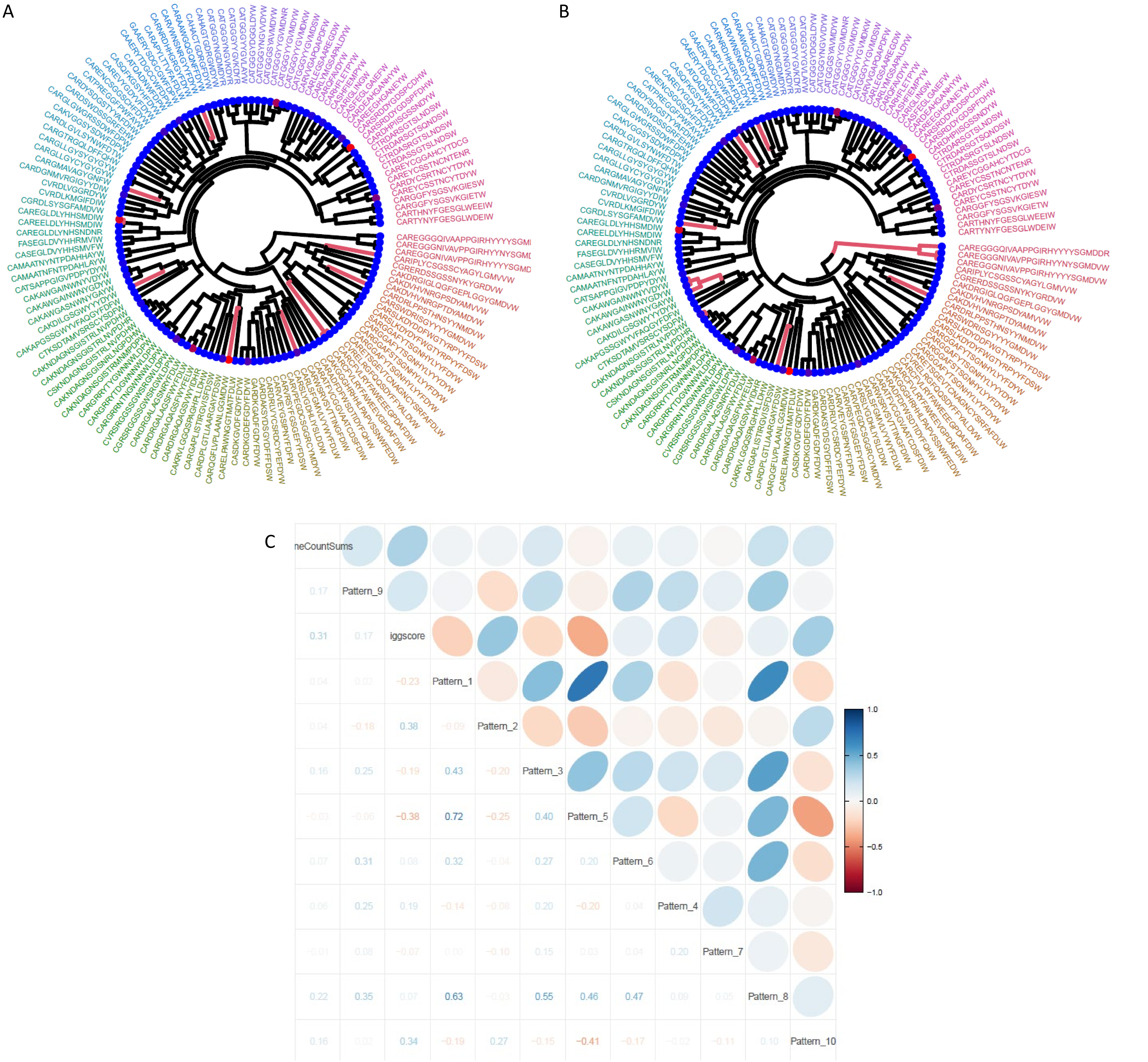

**Supplemental Figure S7.** Alluvial plot illustrating overlap of top 50 expanded IGL (**A**) and IGH (**B**) chains across dataset.

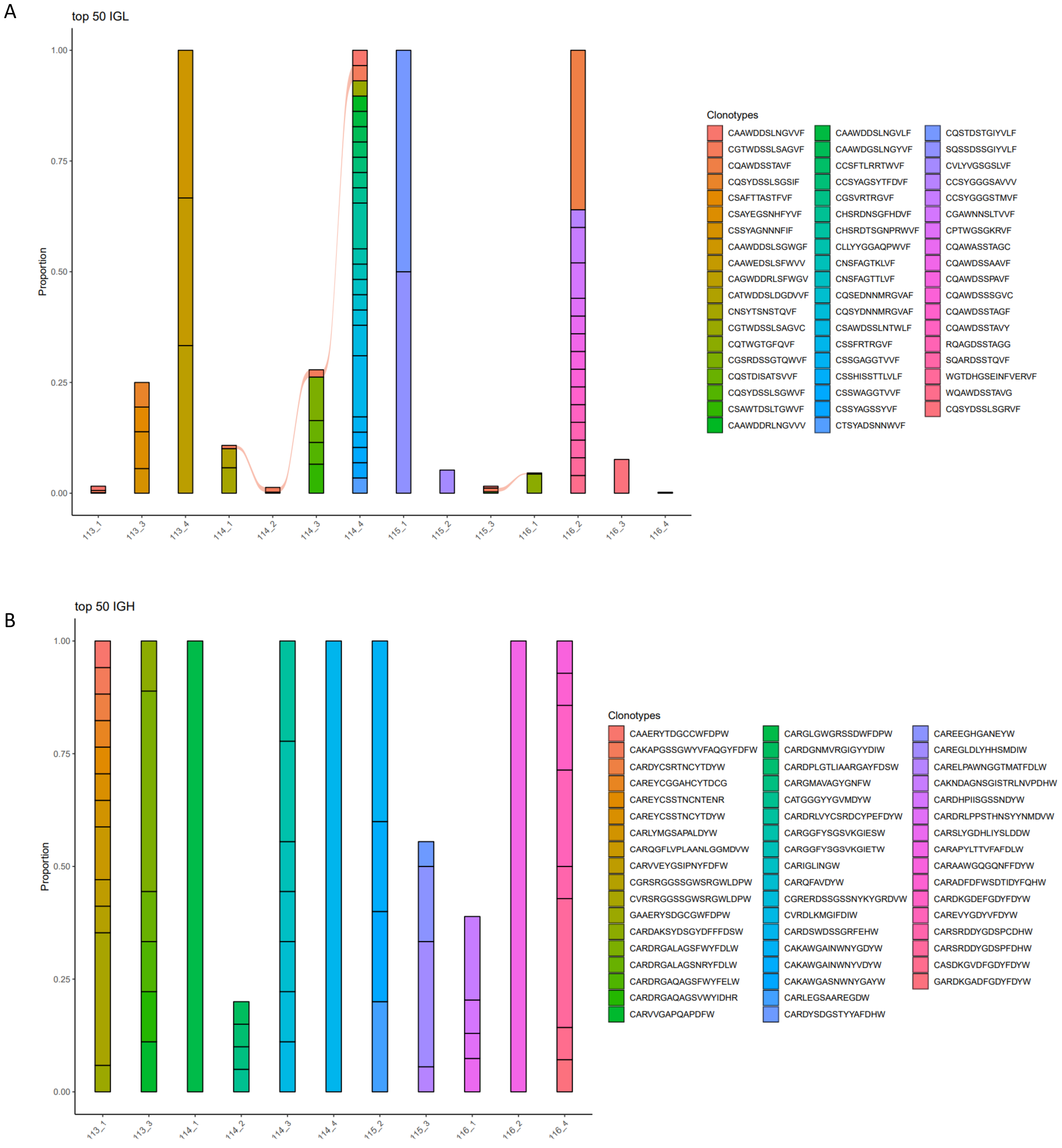

**Supplemental Figure S8.** Spatial BCRs in sample with high TLS density. **A**. Spatial plot showing distribution of top IGH chains. **B**, Heatmap illustrating top IGH chains, chain expression counts and spot annotations.

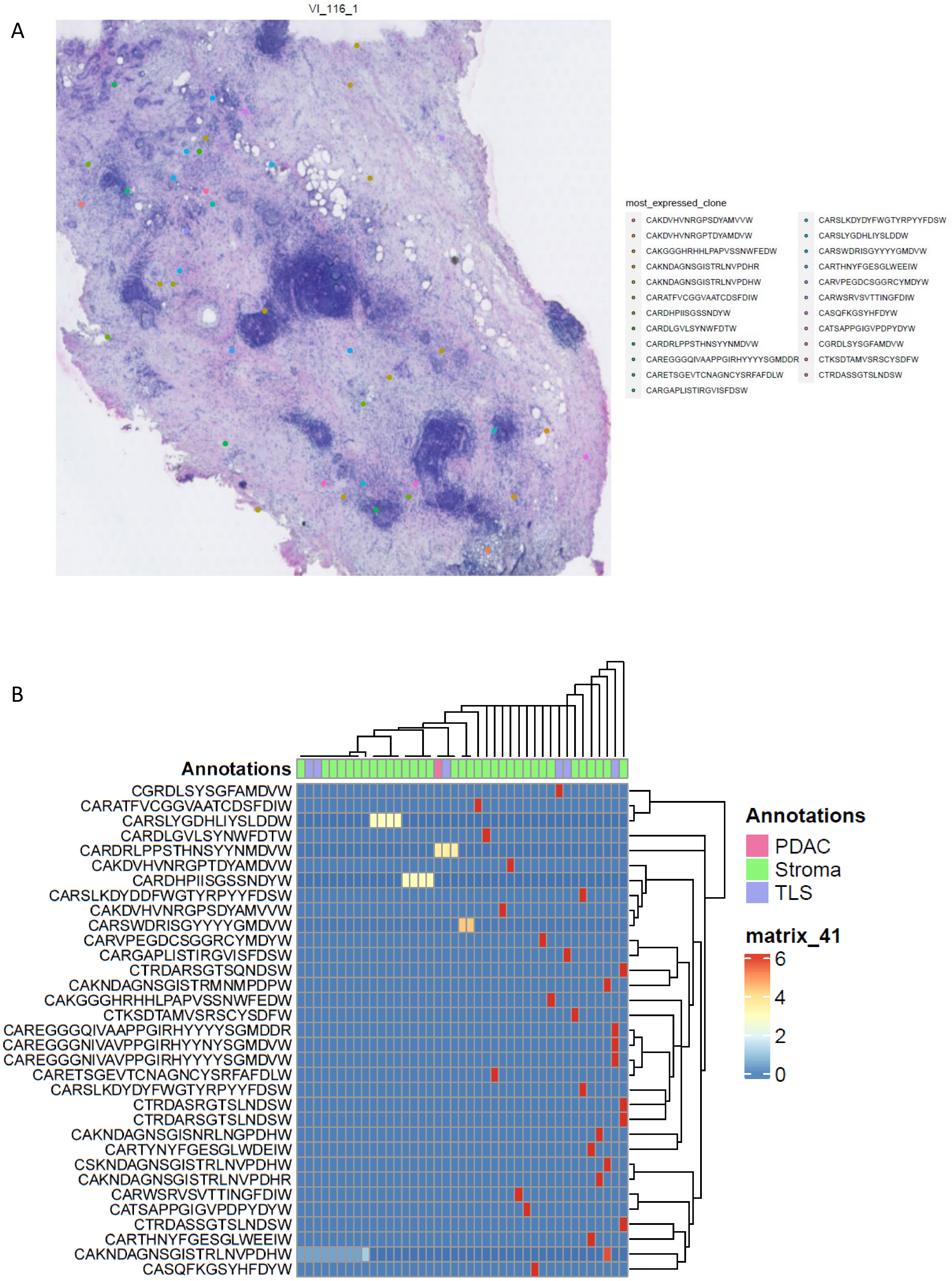

**Supplemental Methods Table S1.** List of antibody-metal conjugates used in imaging mass cytometry

| Channel | Antigen | Clone | Dilution |
|---|---|---|---|
| 89 | **CD45** | D9M81 (CST) | 125 |
| 113 | **PNAd** | MECA-79 (Biolegend) | 250 |
| 115 | **AID** | mAID-2 (Thermo) | 125 |
| 141 | **SMA** | 1A4 | 500 |
| 142 | **Podoplanin** | D2-40 (Biolegend) | 125 |
| 143 | **VIM** | D21H3 | 500 |
| 144 | **CD11c** | EP1347Y (CST) | 250 |
| 145 | **CD45RO** | UCHL1 (Biolegend) | 250 |
| 146 | **CXCR3** | | 100 |
| 147 | **CD69** | EPR21814 | 250 |
| 148 | **CK (Pan-Keratin)** | C11 | 125 |
| 149 | **CD25 (IL2R)** | SP176 (Abcam) | 250 |
| 150 | **PDL1** | E1L3N (CST) | 125 |
| 151 | **CXCR5** | 51505 (Novus) | 125 |
| 152 | **DC-LAMP** | 1010E1.01 (Novus) | 125 |
| 153 | **Tox/Tox2** | E6I3Q (CST) | 250 |
| 154 | **CD57** | HNK-1 (CST) | 250 |
| 155 | **FOXP3** | PCH101 | 75 |
| 156 | **CD4** | EPR6855 | 125 |
| 158 | **ICOS** | D1K2T (CST) | 250 |
| 159 | **CD68** | KP1 | 75 |
| 160 | **CD138** | IHC138 (CST) | 125 |
| 161 | **CD20** | H1 | 125 |
| 162 | **CD8** | C8/144B | 250 |
| 163 | **CD21** | Bu32 (Biolegend) | 250 |
| 164 | **BCL-6** | K112-91 (BD Bio) | 250 |
| 165 | **PD1** | EPR4877 | 250 |
| 166 | **CD45RA** | HI100 | 250 |
| 167 | **GZMB** | D6E9W (CST) | 125 |
| 168 | **KI67** | B56 | 250 |
| 169 | **CD23** | MRQ-57 (Cell Marque) | 125 |
| 170 | **CD3** | Polyclonal, C-terminal | 125 |
| 171 | **LAG3** | 17B4 (Novus) | 125 |
| 172 | **CD137 (4-1BB)** | D2Z4Y (CST) | 250 |
| 173 | **DC-SIGN** | DCN46 (IonPath) | 62.5 |
| 174 | **HLADR** | LN3 | 250 |
| 175 | **CD86** | E2G8P (CST) | 125 |
| 176 | **CCR7** | EPR23192-57 (Abcam) | 250 |
| 191 | **DNA1** | | |
| 193 | **DNA2** | | |
| 195 | **PM2** | 1A36 | 250 |
| 196 | **PM3** | 1A37 | 250 |
| 198 | **PM4** | 1A38 | 250 |
